# The potassium channel subunit K_V_1.8 (*Kcna10*) is essential for the distinctive outwardly rectifying conductances of type I and II vestibular hair cells

**DOI:** 10.1101/2023.11.21.563853

**Authors:** Hannah R. Martin, Anna Lysakowski, Ruth Anne Eatock

**Author notes:** Author for correspondence: Ruth Anne Eatock Dept of Neurobiology, J251 University of Chicago Chicago, IL.

## Abstract

In amniotes, head motions and tilt are detected by two types of vestibular hair cells (HCs) with strikingly different morphology and physiology. Mature type I HCs express a large and very unusual potassium conductance, g_K,L_, which activates negative to resting potential, confers very negative resting potentials and low input resistances, and enhances an unusual non-quantal transmission from type I cells onto their calyceal afferent terminals. Following clues pointing to K_V_1.8 (KCNA10) in the Shaker K channel family as a candidate g_K,L_ subunit, we compared whole-cell voltage-dependent currents from utricular hair cells of K_V_1.8-null mice and littermate controls. We found that K_V_1.8 is necessary not just for g_K,L_ but also for fast-inactivating and delayed rectifier currents in type II HCs, which activate positive to resting potential. The distinct properties of the three K_V_1.8-dependent conductances may reflect different mixing with other K_V_ subunits that are reported to be differentially expressed in type I and II HCs. In K_V_1.8-null HCs of both types, residual outwardly rectifying conductances include K_V_7 (KCNQ) channels.

Current clamp records show that in both HC types, K_V_1.8-dependent conductances increase the speed and damping of voltage responses. Features that speed up vestibular receptor potentials and non-quantal afferent transmission may have helped stabilize locomotion as tetrapods moved from water to land.

**Graphical abstract:** 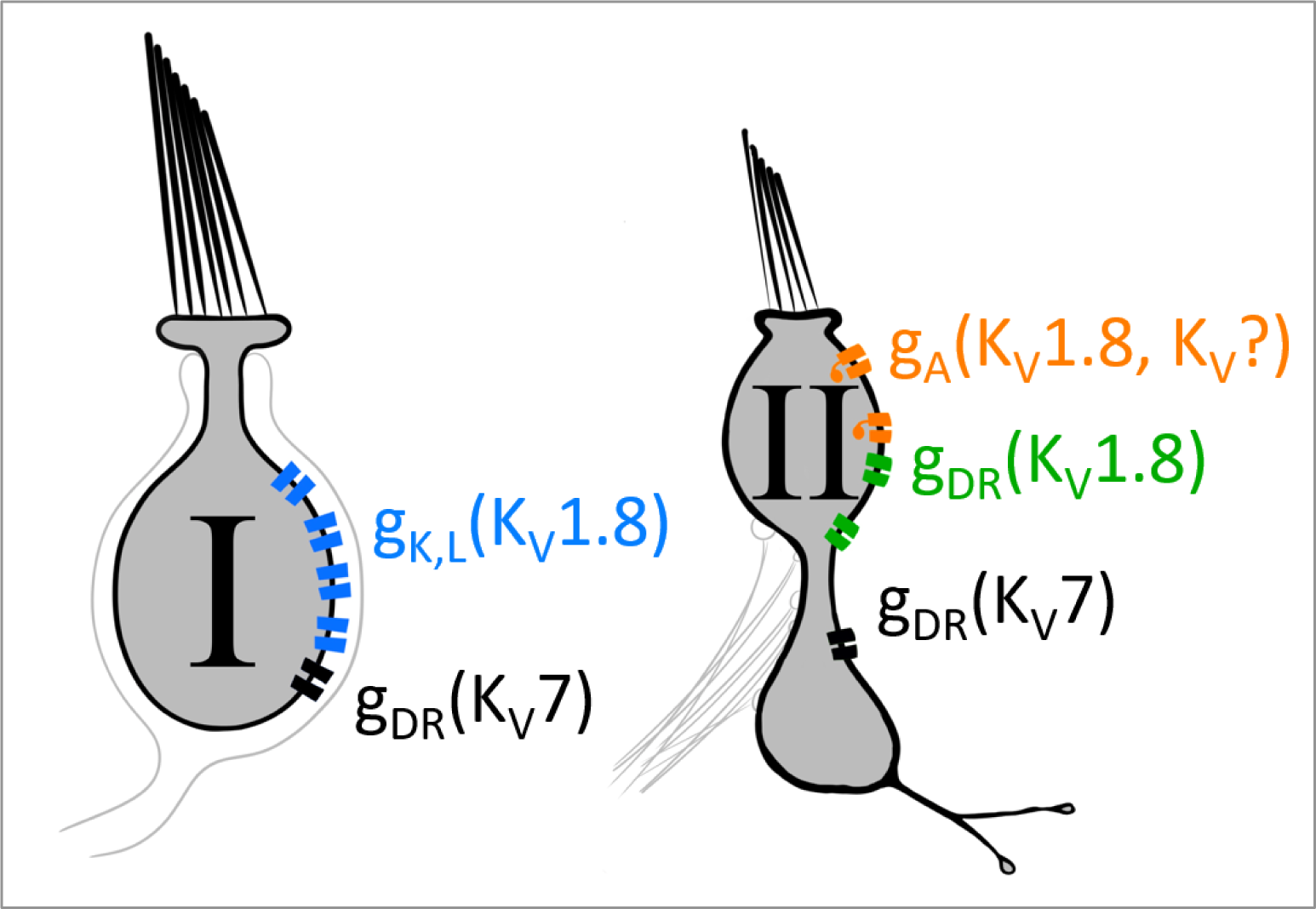

## Introduction

The receptor potentials of hair cells (HCs) are strongly shaped by large outwardly rectifying K^+^ conductances that are differentially expressed according to HC type. Here we report that a specific voltage-gated K^+^ (K_V_) channel subunit participates in very different K_V_ channels dominating the membrane conductances of type I and type II HCs in amniote vestibular organs.

Type I HCs occur only in amniote vestibular organs. Their most distinctive features are that they are enveloped by a calyceal afferent terminal (Wersäll, 1956; Lysakowski and Goldberg, 2004) and that they express g_K,L_ (Correia and Lang, 1990; Rennie and Correia, 1994; Rüsch and Eatock, 1996a): a large non-inactivating conductance with an activation range from –100 to –60 mV, far more negative than other “low-voltage-activated” K_V_ channels. HCs are known for their large outwardly rectifying K+ conductances, which repolarize membrane voltage following a mechanically evoked perturbation and in some cases contribute to sharp electrical tuning of the hair cell membrane. g_K,L_ is unusually large and unusually negatively activated, and therefore strongly attenuates and speeds up the receptor potentials of type I HCs (Correia et al., 1996; Rüsch and Eatock, 1996b). In addition, g_K,L_ augments non-quantal transmission from type I HC to afferent calyx by providing open channels for K^+^ flow into the synaptic cleft (Contini et al., 2012, 2017, 2020; Govindaraju et al., 2023), increasing the speed and linearity of the transmitted signal (Songer and Eatock, 2013).

Type II HCs have compact afferent synaptic contacts (boutons) where the receptor potential drives quantal release of glutamate. They have fast-inactivating (A-type, g_A_) and delayed rectifier (g_DR_) conductances that are opened by depolarization above resting potential (V_rest_).

The unusual properties of g_K,L_ have long attracted curiosity about its molecular nature. g_K,L_ has been proposed to include “M-like” K_V_ channels in the K_V_7 and/or erg channel families (Kharkovets et al., 2000; Hurley et al., 2006; Holt et al., 2007). The K_V_7.4 subunit was of particular interest because it contributes to the low-voltage-activated conductance, g_K,n_, in cochlear outer hair cells, but was eventually eliminated as a g_K,L_ subunit by experiments on K_V_7.4-null mice (Spitzmaul et al., 2013).

Several observations suggested the K_V_1.8 (KCNA10) subunit as an alternative candidate for g_K,L_. K_V_1.8 is highly expressed in vestibular sensory epithelia (Carlisle et al., 2012), particularly hair cells (Lee et al., 2013; Scheffer et al., 2015; McInturff et al., 2018), with slight expression elsewhere (skeletal muscle, Lee et al., 2013; kidney, Yao, 2002). K_V_1.8^−/−^ mice show absent or delayed vestibular-evoked potentials, the synchronized activity of afferent nerve fibers sensitive to fast linear head motions (Lee et al., 2013). Unique among K_V_1 channels, K_V_1.8 has a cyclic nucleotide binding domain (Lang et al., 2000) with the potential to explain g_K,L_’s known cGMP dependence (Behrend et al., 1997; Chen and Eatock, 2000).

Our comparison of whole-cell currents and immunohistochemistry in type I HCs from K_V_1.8^−/−^ and K_V_1.8^+/+,+/−^ mouse utricles confirmed that K_V_1.8 expression is necessary for g_K,L_. More surprisingly, K_V_1.8 expression is also required for A-type and delayed rectifier conductances of type II HCs. In both HC types, eliminating the K_V_1.8-dependent major conductances revealed a smaller delayed rectifier conductance involving K_V_7 channels. Thus, the distinctive outward rectifiers that produce such different receptor potentials in type I and II HCs both include K_V_1.8 and K_V_7 channels.

## Results

We compared whole-cell voltage-activated K^+^ currents in type I and type II hair cells from homozygous knockout (K_V_1.8^−/−^) animals and their wildtype (K_V_1.8^+/+^) or heterozygote (K_V_1.8^+/−^) littermates. We immunolocalized K_V_1.8 subunits in the utricular epithelium and pharmacologically characterized the residual K^+^ currents of K_V_1.8^−/−^ animals. Current clamp experiments demonstrated the impact of K_V_1.8-dependent currents on passive membrane properties.

We recorded from three utricular zones: lateral extrastriola (LES), striola, and medial extrastriola (MES) (Fig. 3A.1); striolar and extrastriolar zones have many structural and functional differences and give rise to afferents with different physiology (reviewed in Goldberg, 2000; Eatock and Songer, 2011). Recordings are from 412 type I and II HCs (53% LES, 30% MES, 17% striola) from mice between postnatal day (P) 5 and P370. We recorded from such a wide age range to test for developmental or senescent changes in the impact of the null mutation. Above P18, we did not see substantial changes in K_V_ channel properties, as reported (González-Garrido et al., 2021).

K_V_1.8^−/−^ animals appeared to be healthy and to develop and age normally, as reported (Lee et al., 2013), and hair cells were healthy (see Methods for criteria).

### K_V_1.8 is necessary for g_K,L_ in type I hair cells

The large low-voltage activated conductance, g_K,L_, in K_V_1.8^+/+,+/−^ type I hair cells produces distinctive whole-cell current responses to voltage steps, as highlighted by our standard type I voltage protocol (Fig. 1A). From a holding potential within the g_K,L_ activation range (here –74 mV), the cell is hyperpolarized to –124 mV, negative to E_K_ and the activation range, producing a large inward current through open g_K,L_ channels that rapidly decays as the channels deactivate. We use the large transient inward current as a hallmark of g_K,L_. The hyperpolarization closes all channels, and then the activation function is probed with a series of depolarizing steps, obtaining the maximum conductance from the peak tail current at –44 mV (Fig. 1A). We detected no difference between the Boltzmann parameters of g_K,L_ G-V curves from K_V_1.8^+/−^ and K_V_1.8^+/+^ type I HCs.

**Figure 1.**
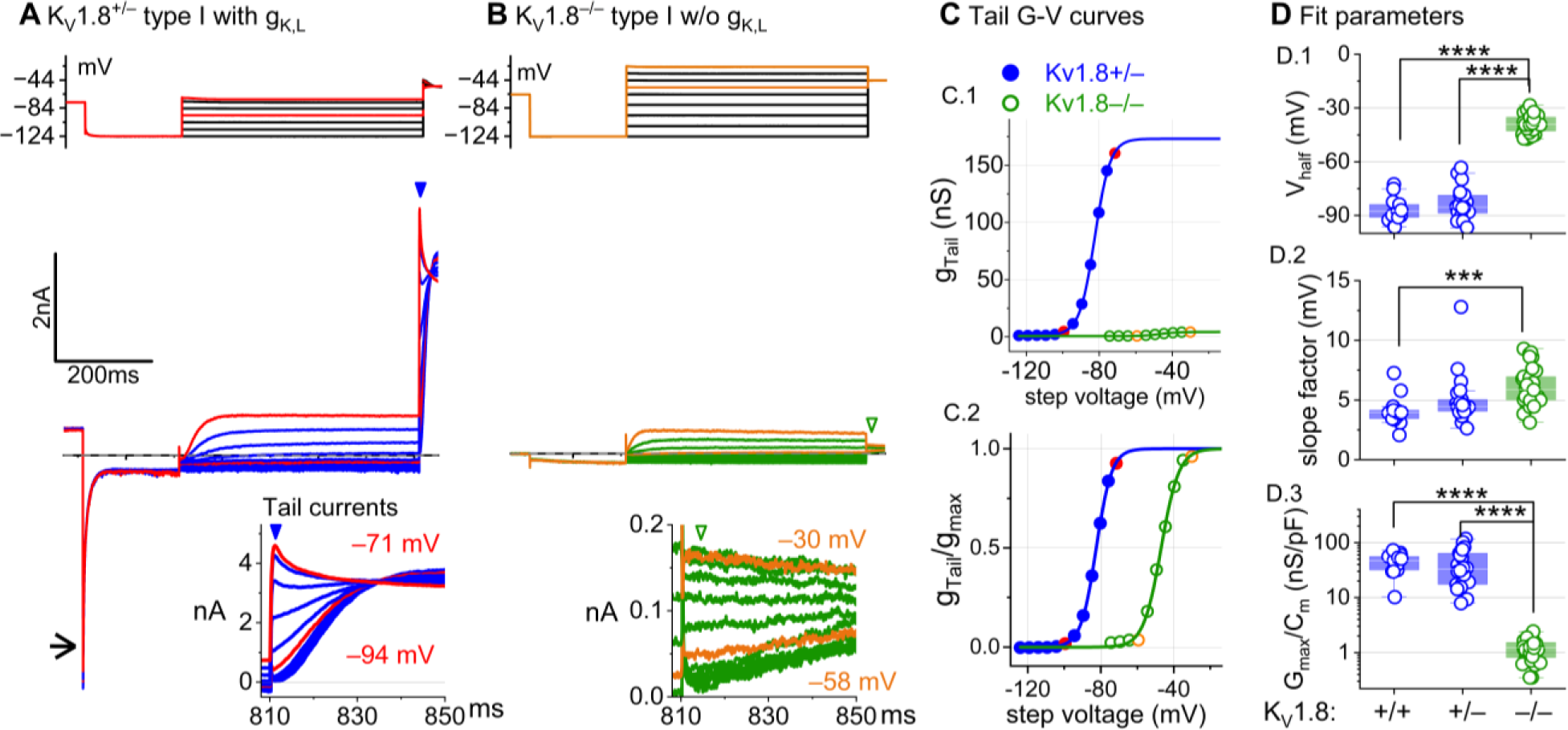
K_V_1.8^−/−^ type I hair cells lacked g_K,L_, the dominant conductance in mature K_V_1.8^+/+,+/−^ type I HCs. Representative voltage-evoked currents in **(A)** a P22 K_V_1.8^+/−^ type I HC and **(B)** a P29 K_V_1.8^−/−^ type I HC. (A) *Arrow,* transient inward current that is a hallmark of g_K,L_. *Arrowheads,* tail currents, magnified in *insets*. For steps positive to the midpoint voltage, tail currents are very large. As a result, K^+^ accumulation in the calyceal cleft reduces driving force on K^+^, causing currents to decay rapidly, as seen in A (Lim et al., 2011). Note that the voltage protocol (top) in B extends to more positive voltages. **(C)** Activation (G-V) curves from tail currents in A and B; symbols, data; curves, Boltzmann fits (Eq. 1). **(D)** Fit parameters from mice >P12 show big effect of K_V_1.8^−/−^ and no difference between K_V_1.8^+/−^ and K_V_1.8^+/+^. *Asterisks* (here and elsewhere): *, p<0.05; **, p<0.01; ***, p<0.001; and ****, p<0.0001. *Line,* median; *Box,* interquartile range; *Whiskers*, outliers. See Table 1 for statistics.

**Table 1.** Type I hair cell K_V_ activation voltage dependence and kinetics. Mean ± SEM (number of cells). g is effect size, Hedge’s g. KWA is Kruskal-Wallis ANOVA.

| Zone | $K_V1.8$ | Tail $V_{1/2}$, mV <sup>a</sup> | Tail $S$, mV <sup>b</sup> | Tail $G_{max}$, nS <sup>c</sup> | Tail $G_{max}/C_m$, nS/pF <sup>d</sup> | Age (median, range) |
| --- | --- | --- | --- | --- | --- | --- |
| Extrastriola | +/+ | $-85 \pm 2$ (12) | $4.3 \pm 0.4$ (12) | $270 \pm 40$ (11) | $47 \pm 8$ (11) | 22, 14-287 |
| | +/- | $-83 \pm 1$ (40) | $5.2 \pm 0.3$ (40) | $210 \pm 20$ (40) | $37 \pm 4$ (40) | 19, 13-259 |
| | -/- | $-40.2 \pm 0.9$ (26) | $5.7 \pm 0.3$ (26) | $5.4 \pm 0.3$ (26) | $1.11 \pm 0.08$ (26) | 45, 14-277 |
| Striola | +/+ | $-87 \pm 3$ (6) | $4.3 \pm 0.3$ (6) | $310 \pm 70$ (6) | $41 \pm 7$ (6) | 40, 15-59 |
| | +/- | $-88 \pm 2$ (3) | $4.7 \pm 0.9$ (3) | $270 \pm 60$ (3) | $44 \pm 6$ (3) | 19, 14-20 |
| | -/- | $-38 \pm 1$ (13) | $6.2 \pm 0.4$ (13) | $6.5 \pm 0.6$ (13) | $1.5 \pm 0.1$ (13) | 202, 14-370 |

<sup>a</sup> -/- vs +/+ : 2-way ANOVA, $p < 1E-9$ , $g$ 7.7; -/- vs +/- : 2-way ANOVA, $p < 1E-9$ , $g$ 6.8
<sup>b</sup> -/- vs +/+ : 2-way ANOVA, $p = 8.4E-4$ , $g$ 1.2
<sup>c</sup> -/- vs +/+ : 2-way ANOVA, $p < 1E-9$ , $g$ 3.7; -/- vs +/- : 2-way ANOVA, $p < 1E-9$ , $g$ 2.1
<sup>d</sup> -/- vs +/+ : 2-way ANOVA, $p < 1E-9$ , $g$ 3.6; -/- vs +/- : 2-way ANOVA, $p < 1E-9$ , $g$ 2.0

For a similar voltage protocol, K_V_1.8^−/−^ type I HCs (Fig. 1B) produced no inward transient current at the step from –74 mV to –124 mV and much smaller depolarization-activated currents during the iterated steps, even at much more positive potentials. Figure 1C compares the conductance-voltage (G-V, activation) curves fit to tail currents (Eq. 1; see insets in Fig. 1A-B): the maximal conductance (g_max_) of the K_V_1.8^−/−^ HC was over 10-fold smaller (Fig. 1C.1), and the curve was positively shifted by >40 mV (Fig. 1C.2). Figure 1D shows the G-V Boltzmann fit parameters for type I HCs from mice >P12, an age at which type I HCs normally express g_K,L_ (Rüsch et al., 1998).

Suppl. Fig. 1 shows how G-V parameters of outwardly rectifying currents in type I HCs changed from P5 to P360. In K_V_1.8^+/+,+,−^ mice, the parameters transitioned over the first 15-20 postnatal days from values for a conventional delayed rectifier, activating positive to resting potential, to g_K,L_ values, as previously described (Rüsch et al., 1998; Géléoc et al., 2004; Hurley et al., 2006). Between P5 and P10, some type I HCs have not yet acquired the physiologically defined conductance, g_K,L_. No effects of K_V_1.8 deletion were seen in the delayed rectifier currents of immature type I HCs (Suppl. Fig. 1B), showing that they were not immature forms of the K_V_1.8-dependent g_K,L_ channels.

K_V_1.8^−/−^ type I HCs had a much smaller residual delayed rectifier that activated positive to resting potential, with V_half_ ∼–40 mV and g_max_ density of 1.3 nS/pF. No additional K^+^ conductance activated up to +40 mV, and G-V parameters did not change much with age from P5 to P370. We characterize this K_V_1.8-independent delayed rectifier later. A much larger non-g_K,L_ delayed rectifier conductance (“g_DR,I_”) was reported in our earlier publication on mouse utricular type I HCs (Rüsch et al., 1998). This current was identified as that remaining after “blocking” g_K,L_ with 20 mM external Ba^2+^. Our new data suggest that there is no large non-g_K,L_ conductance, and that instead high Ba^2+^ positively shifted the apparent voltage dependence of g_K,L_.

### K_V_1.8 strongly affects type I passive properties and responses to current steps

While the cells of K_V_1.8^−/−^ and K_V_1.8^+/−^ epithelia appeared healthy, type I hair cells had smaller membrane capacitances (C_m_): 4-5 pF in K_V_1.8^−/−^ type I HCs, ∼20% smaller than K_V_1.8^+/−^ type I HCs (∼6 pF) and ∼30% smaller than K_V_1.8^+/+^ type I HCs (6-7 pF; Table 2). C_m_ scales with surface area, but soma sizes were unchanged by deletion of K_V_1.8 (Suppl. Table 2). Instead, C_m_ may be higher in K_V_1.8^+/+^ cells because of g_K,L_ for two reasons. First, highly expressed trans-membrane proteins (see discussion of g_K,L_ channel density in Chen and Eatock, 2000) can affect membrane thickness (Mitra et al., 2004), which is inversely proportional to specific C_m_. Second, g_K,L_ could contaminate estimations of capacitive current, which is calculated from the decay time constant of transient current evoked by a small voltage step *outside* the operating range of any ion channels. g_K,L_ has such a negative operating range that, even for V_m_ negative to –90 mV, some g_K,L_ channels are voltage-sensitive and could add to capacitive current.

**Table 2.** Type I hair cell passive membrane properties. Mean ± SEM (number of cells). g is effect size, Hedge’s g. KWA is Kruskal-Wallis ANOVA.

| Zone | $K_V1.8$ | $V_{rest}$, mV <sup>a, b</sup> | $R_{input}$, MΩ <sup>c</sup> | $\tau_{RC}$, ms <sup>d</sup> | $C_m$ (pF) <sup>e</sup> | Age (median, range) |
| --- | --- | --- | --- | --- | --- | --- |
| Extrastriola | +/+ | $-84 \pm 3$ (6) | $44 \pm 6$ (6) | $0.24 \pm 0.03$ (6) | $6.1 \pm 0.4$ (13) | 19.5, 14-287 |
| | +/- | $-88.0 \pm 0.7$ (28) | $55 \pm 5$ (24) | $0.32 \pm 0.03$ (23) | $5.8 \pm 0.2$ (44) | 21, 16-29 |
| | -/- | $-63 \pm 2$ (15) | $1400 \pm 100$ (15) | $6.4 \pm 0.6$ (15) | $5 \pm 0.2$ (27) | 45, 14-202 |
| Striola | +/+ | $-87 \pm 2$ (4) | $50 \pm 8$ (4) | $0.30 \pm 0.04$ (4) | $7.4 \pm 0.7$ (7) | 43, 40-59 |
| | +/- | $-87 \pm 3$ (3) | $38 \pm 8$ (2) | $0.21 \pm 0.01$ (2) | $5.9 \pm 0.6$ (3) | 19, 19-20 |
| | -/- | $-74 \pm 5$ (5) | $1000 \pm 300$ (4) | $4.2 \pm 1.0$ (4) | $4.4 \pm 0.2$ (14) | 202, 24-370 |

<sup>a</sup> striolar -/- vs ES -/- : 2-way ANOVA, $p = 0.006$ , $g$ 1.2; striolar -/- vs striolar +/+, +/- : 2-way ANOVA, $p = 0.005$ , $g$ 1.7
<sup>b</sup> -/- vs +/+ : 2-way ANOVA, $p < 1E-9$ , $g$ 2.3; -/- vs +/- : 2-way ANOVA, $p < 1E-9$ , $g$ 3.4
<sup>c</sup> -/- vs +/+ : 2-way ANOVA, $p < 1E-9$ , $g$ 3.1; -/- vs +/- : 2-way ANOVA, $p < 1E-9$ , $g$ 3.9
<sup>d</sup> -/- vs +/+ : 2-way ANOVA, $p < 1E-9$ , $g$ 2.7; -/- vs +/- : 2-way ANOVA, $p < 1E-9$ , $g$ 3.4
<sup>e</sup> -/- vs +/+ : 2-way ANOVA, $p = 3E-7$ , $g$ 1.5; -/- vs +/- : 2-way ANOVA, $p = 1.3E-4$ , $g$ 1.0; +/- vs +/+ : 2-way ANOVA, $p = 0.048$ , $g$ 0.6

Basolateral conductances help set the resting potential and passive membrane properties that regulate the time course and gain of voltage responses to small currents. To examine the effect of K_V_1.8 on these properties, we switched to current clamp mode and measured resting potential (V_rest_), input resistance (R_in_, equivalent to voltage gain for small current steps, △V/△I), and membrane time constant (τ_RC_). In K_V_1.8^−/−^ type I HCs, V_rest_ was much less negative (Fig. 2A.1), R_in_ was greater by ∼20-fold (Fig. 2A.2), and membrane charging times were commensurately longer (Fig. 2A.3).

**Figure 2.**
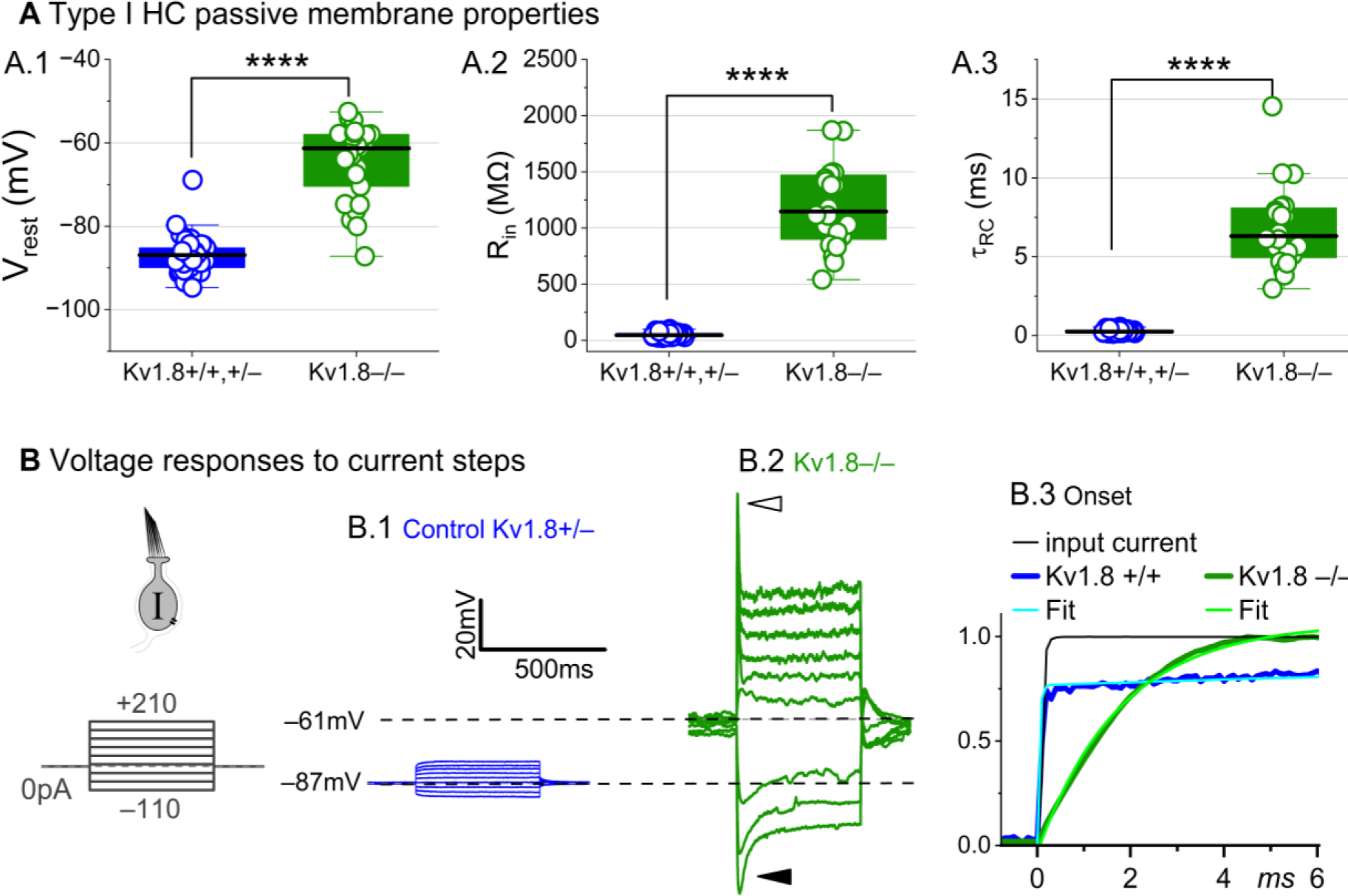
K_V_1.8^−/−^ type I hair cells had much longer membrane charging times and higher input resistances (voltage gains) than K_V_1.8^+/+,+/−^ type I HCs. **(A)** g_K,L_ strongly affects passive membrane properties: (**A.1**) V_rest_, (**A.2**) R_in_, input resistance, and (**A.3**) membrane time constant, τ_RC_ = (R_input_ * C_m_). See Table 2 for statistics. **(B)** Current clamp responses to the same scale from (B.1) K_V_1.8^+/−^ and (B.2) K_V_1.8^−/−^ type I cells, both P29. *Filled arrowhead (B.2),* sag indicating I_H_ activation. *Open arrowhead,* Depolarization rapidly decays as I_DR_ activates. B.3, The 1^st^ 6 ms of voltage responses to 170-pA injection is normalized to steady-state value; overlaid curves are double-exponential fits (K_V_1.8^+/+^, τ 40 ps and 2.4 ms) and single-exponential fits (K_V_1.8^−/−^, τ 1.1 ms).

The differences between the voltage responses of K_V_1.8^+/+,+/−^ and K_V_1.8^−/−^ type I HCs are expected from the known impact of g_K,L_ on V_rest_ and R_in_ (Correia and Lang, 1990; Ricci et al., 1996; Rüsch and Eatock, 1996b; Songer and Eatock, 2013). The large K^+^-selective conductance at V_rest_ holds V_rest_ close to E_K_ (K^+^ equilibrium potential) and minimizes gain (△V/△I), such that voltage-gated conductances are negligibly affected by the input current and the cell produces approximately linear, static responses to iterated current steps. For K_V_1.8^−/−^ type I HCs, with their less negative V_rest_ and larger R_in_, positive current steps evoked a fast initial depolarization (Fig. 2B.2), activating residual delayed rectifiers and repolarizing the membrane toward E_K_. Negative current steps evoked an initial “sag” (Fig. 2B.2), a hyperpolarization followed by slow repolarization as the HCN1 channels open (Rüsch and Eatock, 1996b; Horwitz et al., 2011).

Overall, comparison of the K_V_1.8^+/+,+/−^ and K_V_1.8^−/−^ responses shows that with K_V_1.8 (g_K,L_), the voltage response of the type I hair cell is smaller but better reproduces the time course of the input current.

### KV1.8 is necessary for both inactivating and non-inactivating K_V_ currents in type II hair cells

Type II HCs also express K_V_1.8 mRNA (McInturff et al., 2018; Orvis et al., 2021). Although their outwardly-rectifying conductances (g_A_ and g_DR_) differ substantially in voltage dependence and size from g_K,L_, both conductances were strongly affected by the null mutation: g_A_ was eliminated and the delayed rectifier was substantially smaller. Below we describe g_A_ and g_DR_ in K_V_1.8^+/+,+/−^ type II HCs and the residual outward rectifying current in K_V_1.8^−/−^ type II HCs.

***K_V_1.8^+/+,+/−^ type II HCs.*** Most (81/84) K_V_1.8^+/+,+/−^ type II HCs expressed a rapidly-activating, rapidly-inactivating A-type conductance (g_A_). We define A current as the outwardly rectifying current that inactivates by over 30% within 200 ms. g_A_ was more prominent in extrastriolar zones, as reported (Holt et al. 1999, Weng and Correia 1999).

We compared the activation and inactivation time course and inactivation prominence for 200-ms steps from –124 mV to ∼30 mV. Outward currents fit with Eq. 3 yielded fast inactivation time constants (τ_Inact, Fast_) of ∼30 ms in LES (Fig. 3A.2). τ_Inact, Fast_ was faster in LES than in MES or striola (Fig. 3A.3) and fast inactivation was a larger fraction of the total inactivation in LES than striola (∼0.5 vs. 0.3, Fig. 3A.4).

**Figure 3.**
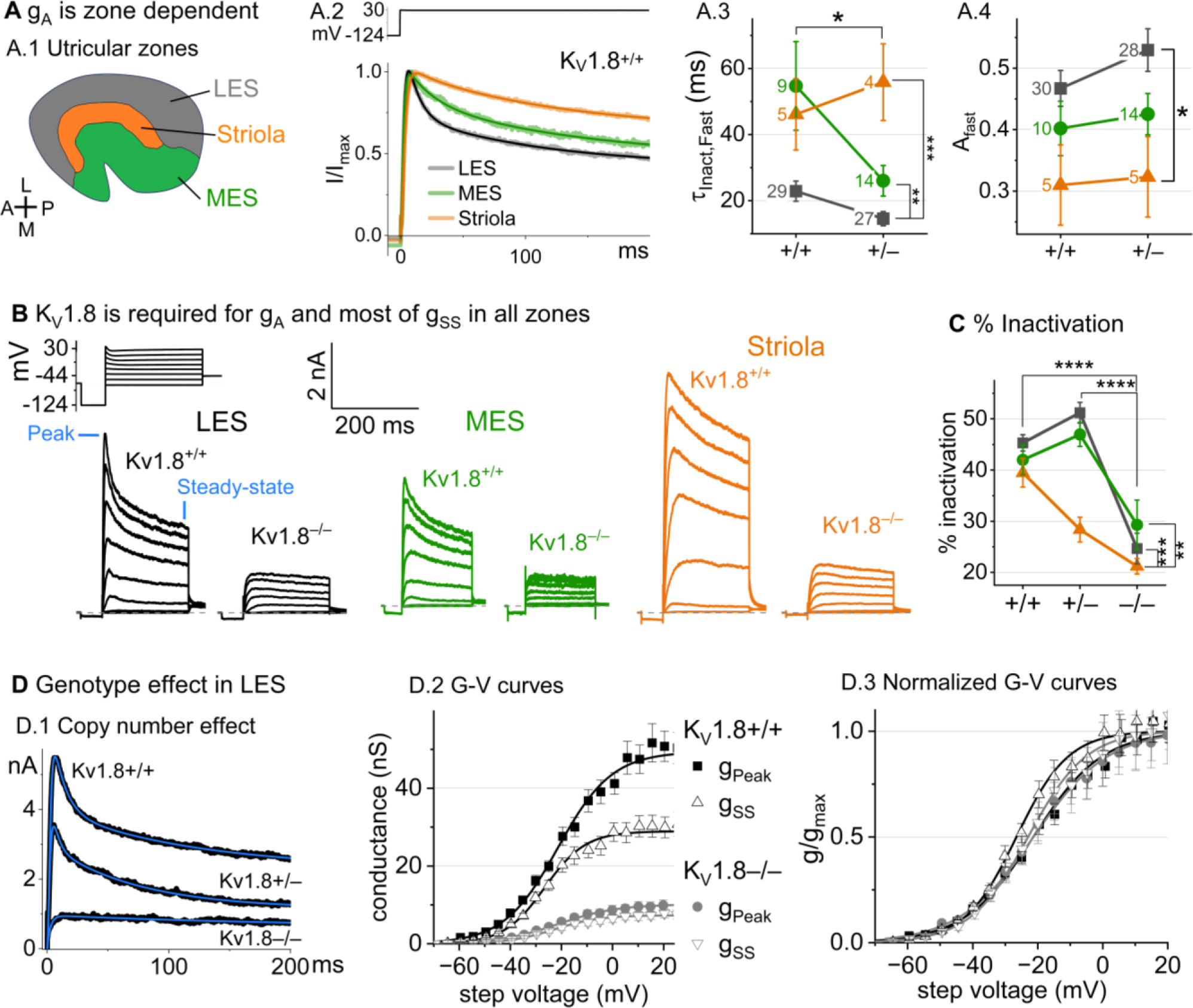
KV1.8^−/−^ type II HCs in all zones of the sensory epithelium lacked the major rapidly inactivating conductance, gA, and had less delayed rectifier conductance. Activation and inactivation varied with epithelial zone and genotype. (**A**) gA inactivation time course varied across zones. (**A.1**) Zones of the utricular epithelium. (**A.2**) Normalized currents evoked by steps from –124 mV to +30 mV with overlaid fits of Eq. 3. (**A.3**) tInact,Fast was faster in KV1.8^+/−^ than KV1.8^+/+^ HCs, and faster in LES than other zones. Brackets show post hoc pairwise comparisons between two zones (vertical brackets) and horizontal brackets compare two genotypes; see Table 3 for statistics. (**A.4**) Fast inactivation was a greater fraction of total inactivation in LES than striola. (B) Exemplars; ages, *left to right*, P312, P53, P287, P49, P40, P154. (C) % inactivation at 30 mV was much lower in KV1.8^−/−^ than KV1.8^+/−^ and KV1.8^+/+^, and lower in striola than LES and MES. Interaction between zone and genotype was significant (Table 3). (D) Exemplar currents and G-V curves from LES type II HCs show a copy number effect. (**D.1**) Currents for examples of the 3 genotypes evoked by steps from –124 mV to +30 mV fit with Eq. 3. (**D.2**) Averaged peak and steady-state conductance-voltage datapoints from LES cells (+/+, n=37; –/–, n=20) were fit with Boltzmann equations (Eq. 1) and normalized by gmax in (**D.3**). See Table 4 for statistics.

To show voltage dependence of activation, we generated G-V curves for peak currents (sum of A-current and delayed rectifier) and steady-state currents measured at 200 ms, after g_A_ has mostly inactivated (Figure 3D.2). K_V_1.8^+/−^ HCs had smaller currents than K_V_1.8^+/+^ HCs, reflecting a smaller g_DR_ (Fig. 3D) and faster fast inactivation (Fig. 3A.3). As discussed later, these effects may relate to effects of the K_V_1.8 gene dosage on the relative numbers of different K_V_1.8 heteromers.

For K_V_1.8^+/+^ and K_V_1.8^+/−^ HCs, the voltage dependence as summarized by V_half_ and slope factor (S) was similar. Relative to g_SS_, g_Peak_ had a more positive V_half_ (∼–21 vs. ∼–26) and greater S (∼12 vs. ∼9, Fig. 3D, Table 4). Because g_Peak_ includes channels with and without fast inactivation, the shallower g_Peak_-V curve may reflect a more heterogeneous channel population. Only g_Peak_ showed zonal variation, with more positive V_half_ in LES than striola (∼–20 mV vs. ∼–24 mV, Fig. 3D, Table 4). We later suggest that variable subunit composition may drive zonal variation in g_Peak_.

**Table 3.**
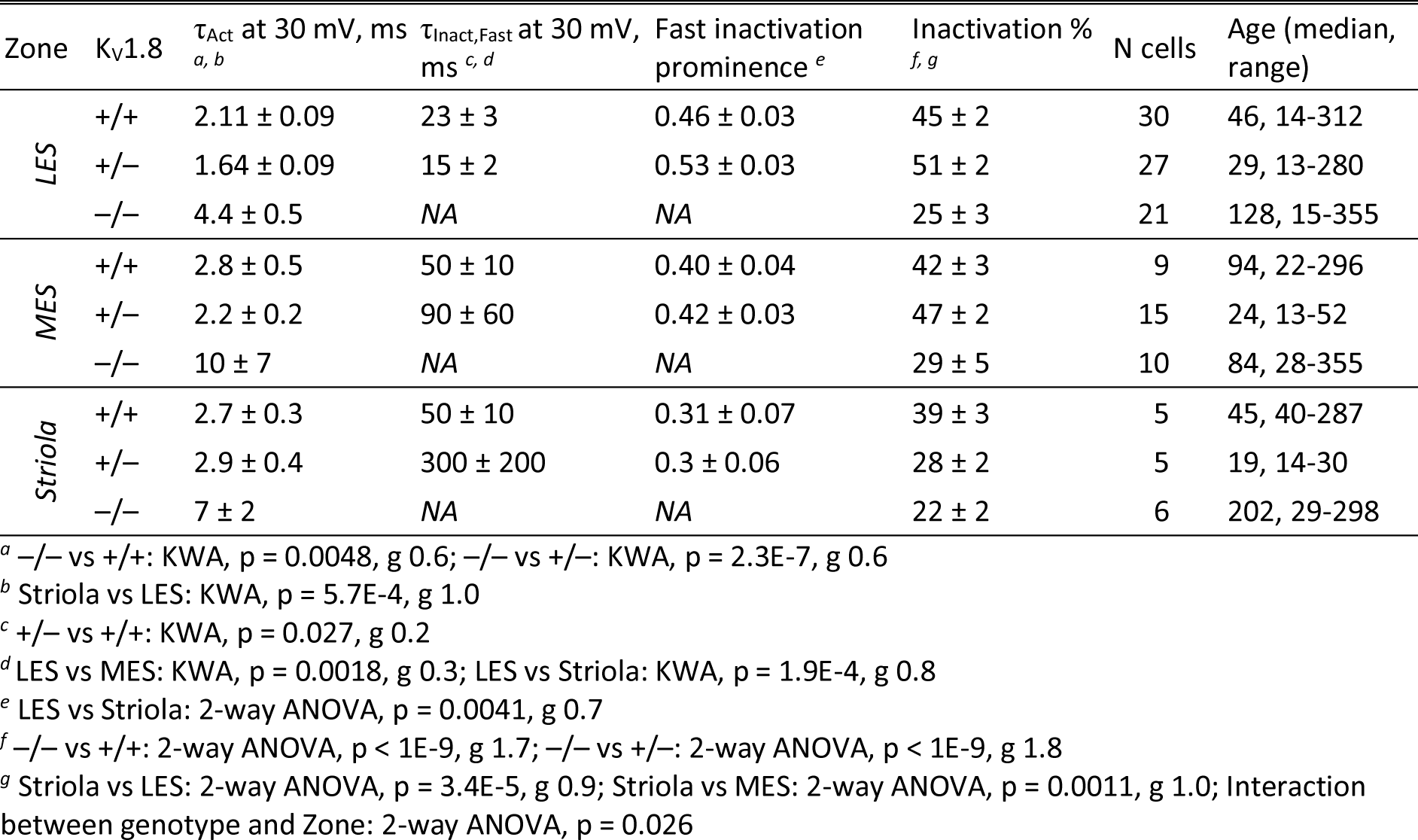
Type II hair cell K_V_ currents: Activation and inactivation time course at +30 mV. Mean ± SEM. g is effect size, Hedge’s g. KWA is Kruskal-Wallis ANOVA.

| Zone | $K_V1.8$ | $\tau_{Act}$ at 30 mV, ms<br>*a, b* | $\tau_{Inact, Fast}$ at 30 mV, ms<br>*c, d* | Fast inactivation prominence<br>*e* | Inactivation %<br>*f, g* | N cells | Age (median, range) |
| --- | --- | --- | --- | --- | --- | --- | --- |
| LES | +/+ | 2.11 ± 0.09 | 23 ± 3 | 0.46 ± 0.03 | 45 ± 2 | 30 | 46, 14-312 |
| | +/- | 1.64 ± 0.09 | 15 ± 2 | 0.53 ± 0.03 | 51 ± 2 | 27 | 29, 13-280 |
| | -/- | 4.4 ± 0.5 | NA | NA | 25 ± 3 | 21 | 128, 15-355 |
| MES | +/+ | 2.8 ± 0.5 | 50 ± 10 | 0.40 ± 0.04 | 42 ± 3 | 9 | 94, 22-296 |
| | +/- | 2.2 ± 0.2 | 90 ± 60 | 0.42 ± 0.03 | 47 ± 2 | 15 | 24, 13-52 |
| | -/- | 10 ± 7 | NA | NA | 29 ± 5 | 10 | 84, 28-355 |
| Striola | +/+ | 2.7 ± 0.3 | 50 ± 10 | 0.31 ± 0.07 | 39 ± 3 | 5 | 45, 40-287 |
| | +/- | 2.9 ± 0.4 | 300 ± 200 | 0.3 ± 0.06 | 28 ± 2 | 5 | 19, 14-30 |
| | -/- | 7 ± 2 | NA | NA | 22 ± 2 | 6 | 202, 29-298 |

<sup>a</sup> -/- vs +/+ : KWA, $p = 0.0048$ , $g = 0.6$ ; -/- vs +/- : KWA, $p = 2.3E-7$ , $g = 0.6$
<sup>b</sup> Striola vs LES: KWA, $p = 5.7E-4$ , $g = 1.0$
<sup>c</sup> +/- vs +/+ : KWA, $p = 0.027$ , $g = 0.2$
<sup>d</sup> LES vs MES: KWA, $p = 0.0018$ , $g = 0.3$ ; LES vs Striola: KWA, $p = 1.9E-4$ , $g = 0.8$
<sup>e</sup> LES vs Striola: 2-way ANOVA, $p = 0.0041$ , $g = 0.7$
<sup>f</sup> -/- vs +/+ : 2-way ANOVA, $p < 1E-9$ , $g = 1.7$ ; -/- vs +/- : 2-way ANOVA, $p < 1E-9$ , $g = 1.8$
<sup>g</sup> Striola vs LES: 2-way ANOVA, $p = 3.4E-5$ , $g = 0.9$ ; Striola vs MES: 2-way ANOVA, $p = 0.0011$ , $g = 1.0$ ; Interaction between genotype and Zone: 2-way ANOVA, $p = 0.026$

**Table 4.** Type II hair cell K_V_ currents: Activation voltage dependence. Mean ± SEM. g is effect size, Hedge’s g. KWA is Kruskal-Wallis ANOVA.

| Zone | $K_V1.8$ | Peak $V_{1/2}$, mV<br>*a* | Peak $S$, mV<br>*b, c* | A-type $g_{max}/C_m$, nS/pF<br>*d* | SS $V_{half}$, mV<br>*e* | SS $S$, mV<br>*f* | SS $g_{max}/C_m$, nS/pF<br>*g, h* | N cells | Age (median, range) |
| --- | --- | --- | --- | --- | --- | --- | --- | --- | --- |
| LES | +/+ | -19.8 ± 0.6 | 11.8 ± 0.4 | 4.0 ± 0.3 | -25.0 ± 0.5 | 8.7 ± 0.3 | 7.1 ± 0.8 | 37 | 46, 14-312 |
| | +/- | -19.8 ± 0.8 | 12.8 ± 0.4 | 3.8 ± 0.3 | -26.8 ± 0.8 | 8.7 ± 0.3 | 4.9 ± 0.4 | 35 | 29, 13-280 |
| | -/- | -18 ± 1 | 11.7 ± 0.4 | 0.37 ± 0.05 | -19 ± 1 | 12.1 ± 0.5 | 1.8 ± 0.2 | 20 | 128, 15-355 |
| MES | +/+ | -22 ± 1 | 11 ± 0.7 | 4.1 ± 0.7 | -26 ± 1 | 8.3 ± 0.5 | 9 ± 1 | 11 | 94, 22-296 |
| | +/- | -21 ± 1 | 11.8 ± 0.4 | 3.6 ± 0.5 | -27 ± 1 | 9.0 ± 0.3 | 5.9 ± 0.7 | 16 | 24, 13-52 |
| | -/- | -19 ± 1 | 10.8 ± 0.6 | 0.6 ± 0.1 | -20 ± 1 | 10.7 ± 0.7 | 2.5 ± 0.3 | 15 | 84, 28-355 |
| Striola | +/+ | -24 ± 1 | 9.6 ± 0.5 | 5 ± 1 | -26.6 ± 0.9 | 8.2 ± 0.4 | 12 ± 1 | 7 | 45, 40-287 |
| | +/- | -25 ± 2 | 9.4 ± 0.4 | 2.6 ± 0.6 | -28 ± 2 | 8.2 ± 0.3 | 10 ± 2 | 6 | 19, 14-30 |
| | -/- | -21.3 ± 0.9 | 10.3 ± 0.5 | 0.7 ± 0.1 | -21.7 ± 0.8 | 10.5 ± 0.6 | 3.9 ± 0.5 | 8 | 202, 29-298 |

<sup>a</sup> Striola vs LES: 2-way ANOVA, $p = 0.00116$ , $g = 0.9$
<sup>b</sup> Striola vs MES: 2-way ANOVA, $p = 0.016$ , $g = 0.8$ ; Striola vs LES: 2-way ANOVA, $p = 7.5E-6$ , $g = 1.2$
<sup>c</sup> -/- vs +/- : 2-way ANOVA, $p = 0.036$ , $g = 0.5$
<sup>d</sup> -/- vs +/+ : Welch ANOVA, $p < 1E-9$ , $g = 2.3$ ; -/- vs +/- : Welch ANOVA, $p < 1E-9$ , $g = 2.3$
<sup>e</sup> -/- vs +/+ : 2-way ANOVA, $p < 1E-9$ , $g = 1.4$ ; -/- vs +/- : 2-way ANOVA, $p < 1E-9$ , $g = 1.6$
<sup>f</sup> -/- vs +/+ : 2-way ANOVA, $p < 1E-9$ , $g = 1.4$ ; -/- vs +/- : 2-way ANOVA, $p = 4.5E-7$ , $g = 1.1$
<sup>g</sup> -/- vs +/+ : Welch ANOVA, $p < 1E-9$ , $g = 1.6$ ; -/- vs +/- : Welch ANOVA, $p < 1E-9$ , $g = 1.3$ ; +/- vs +/- : Welch ANOVA, $p = 0.007$ , $g = 1.6$
<sup>h</sup> Striola vs LES: 1-way ANOVA, $p = 0.001$ , $g = 0.9$ ; Striola vs MES: 1-way ANOVA, $p = 0.01$ , $g = 0.8$

***K_V_1.8^−/−^ type II HCs*** from all zones were missing g_A_ and 30-50% of g_DR_ (Fig. 3B-D). The residual delayed rectifier (1.3 nS/pF) had a more positive V_half_ than g_DR_ in K_V_1.8^+/+,+/−^ HCs (∼–20 mV *vs.* ∼–26 mV, Fig. 3D.2). We refer to the K_V_1.8-dependent delayed rectifier component as g_DR_(K_V_1.8) and to the residual, K_V_1.8-*in*dependent delayed rectifier component as g_DR_(K_V_7) because, as we show later, it includes K_V_7 channels. Supplemental Figure 3A shows the development of K_V_1.8-dependent and independent K_V_ currents in type II HCs with age from P5 to over P300. In K_V_1.8^+/+,+/−^ type II HCs, g_A_ was present at all ages with a higher % inactivation after P18 than at P5-P10 (Suppl. Fig. 3A.4). g_Peak_ did not change much above P12 except for a compression of conductance density from P13 to P370 (partial correlation coefficient = –0.4, p = 2E-5, Suppl. Fig. 3A.3).

We saw small rapidly inactivating outward currents in a minority of K_V_1.8^−/−^ type II HCs (23%, 7/30), all >P12 and extrastriolar (Suppl. Fig. 4). These currents overlapped with g_A_ in percent inactivation, inactivation kinetics, and activation voltage dependence but were very small. As discussed later, we suspect that these currents flow through homomers of inactivating K_V_ subunits that in control hair cells join with K_V_1.8 subunits and confer inactivation on the heteromeric conductance.

### K_V_1.8 affects type II passive properties and responses to current steps

In type II HCs, absence of K_V_1.8 did not change V_rest_ (Fig. 4A.1) because g_A_ and g_DR_ both activate positive to rest, but significantly increased R_in_ and τ_RC_ (Fig. 4A.2-A.3).

**Figure 4.**
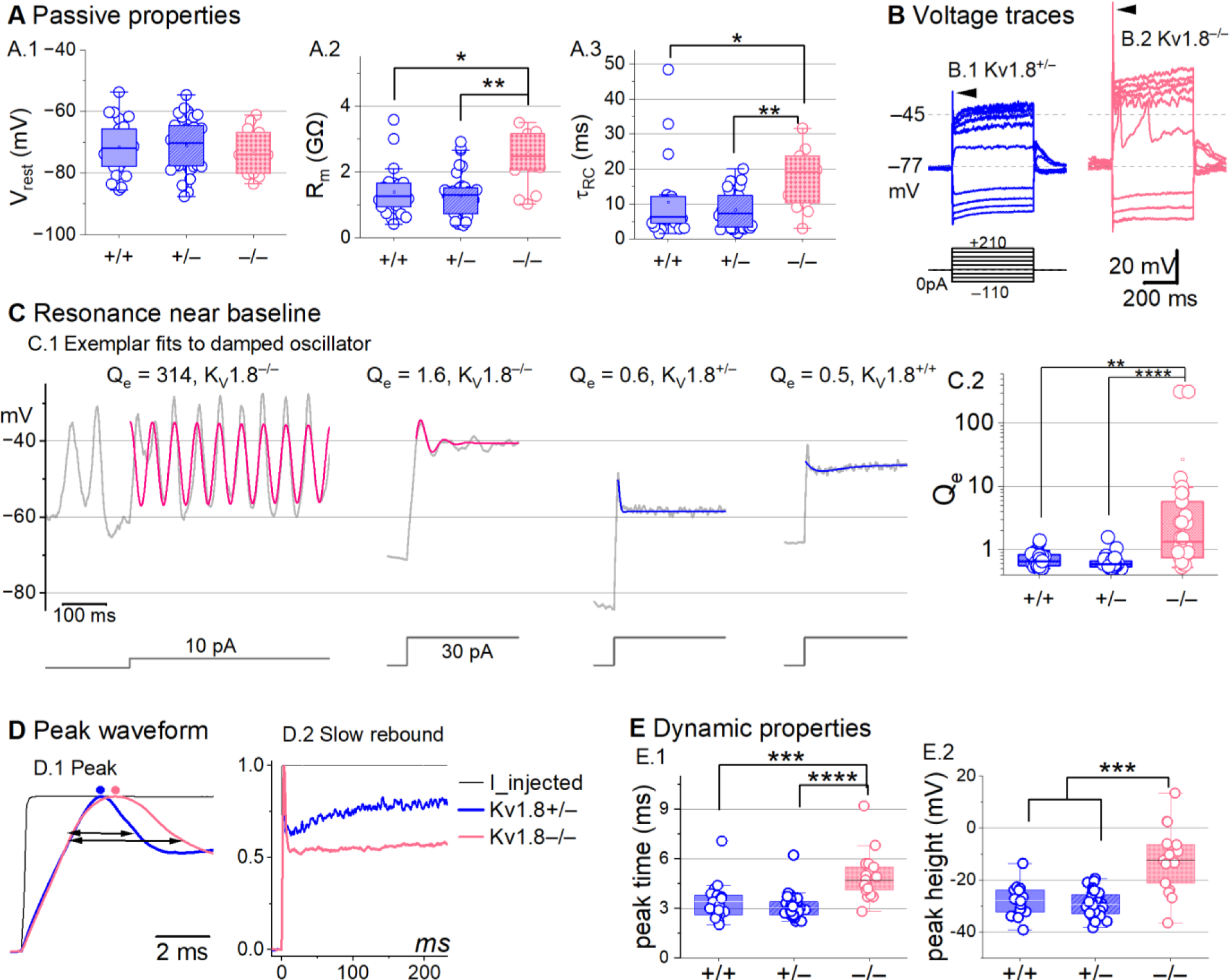
K_V_1.8^−/−^ type II hair cells had larger, slower voltage responses and more electrical resonance. **(A)** Passive membrane properties near resting membrane potential: A.1) Resting potential. R_input_ (A.2) and τ_RC_ (A.3) were obtained from single exponential fits to voltage responses < 15 mV. See Table 5 for statistics. (**B**) Exemplar voltage responses to iterated current steps (*bottom)* illustrate key changes in gain and resonance with K_V_1.8 knockout. (**B.1**) K_V_1.8^+/−^ type II HC (P24, LES) and (**B.2**) K_V_1.8^−/−^ type II HC (P53, LES). *Arrowheads,* depolarizing transients. (**C**) Range of resonance illustrated for K_V_1.8^−/−^ type II HCs (*left, pink curves fit to* *Eq. 5*) and controls (*right, blue fits*). (**C.1**) *Resonant frequencies, left to right:* 19.6, 18.4, 34.4, 0.3 Hz. Leftmost cell resonated spontaneously (before step). (**C.2**) Tuning quality (Q_e_; Eq. 6) was higher for K_V_1.8^−/−^ type II HCs (KWA, p = 0.0064 vs. K_V_1.8^+/+^; p = 7E-8 vs. K_V_1.8^+/−^). (**D**) K_V_1.8^−/−^ type II HCs had higher, slower peaks and much slower rebound potentials in response to large (170-pA) current steps. (**D.1**) Normalized to show initial depolarizing transient (*filled circles*, times of peaks; *horizontal arrows*, peak width at half-maximum). (**D.2**) Longer time scale to highlight how null mutation reduced post-transient rebound. (**E**) In K_V_1.8^−/−^ HCs, depolarizing transients evoked by a +90-pA step were slower to peak (**E.1**) and (**E.2**) larger.

**Table 5.** Type II hair cell passive membrane properties. Mean ± SEM (number of cells). g is effect size, Hedge’s g. KWA is Kruskal-Wallis ANOVA.

| Zone | K<sub>v</sub>1.8 | V<sub>rest</sub>, mV | R<sub>input</sub>, GΩ<sup>a</sup> | τ<sub>RC</sub>, ms<sup>b</sup> | Peak height, mV, 170<sup>c</sup> | Peak time, ms<sup>d</sup> | C<sub>m</sub>, pF | Age (median, range) |
| --- | --- | --- | --- | --- | --- | --- | --- | --- |
| Extrastriola | +/+ | -71 ± 2 (19) | 1.4 ± 0.2 (16) | 11 ± 3 (16) | -20 ± 2 (15) | 2.5 ± 0.2 (15) | 4.7 ± 0.2 (50) | 45, 16-312 |
| | +/- | -71 ± 2 (34) | 1.2 ± 0.1 (27) | 9 ± 1 (27) | -20 ± 1 (30) | 2.44 ± 0.08 (30) | 4.6 ± 0.1 (52) | 27, 13-280 |
| | -/- | -76 ± 2 (9) | 2.3 ± 0.3 (7) | 16 ± 3 (7) | 2 ± 6 (7) | 3.6 ± 0.3 (7) | 4.6 ± 0.2 (35) | 53, 15-154 |
| Striola | +/+ | -73.1 ± 1.0 (6) | 1.4 ± 0.1 (6) | 9 ± 1 (6) | -20 ± 2 (5) | 2.7 ± 0.1 (5) | 4.6 ± 0.2 (7) | 45, 40-224 |
| | +/- | -71 ± 3 (5) | 1.4 ± 0.3 (6) | 7 ± 2 (6) | -20 ± 2 (6) | 2.3 ± 0.1 (6) | 4.8 ± 0.2 (6) | 19, 19-30 |
| | -/- | -68 ± 2 (6) | 3.0 ± 0.7 (6) | 26 ± 10 (6) | 2 ± 7 (4) | 4 ± 1 (4) | 4.4 ± 0.3 (7) | 178, 29-298 |

<sup>a</sup> -/- vs +/+: KWA, p = 0.015, g 1.2; -/- vs +/-: KWA, p = 0.002, g 1.5
<sup>b</sup> -/- vs +/+: KWA, p = 0.016, g 0.7; -/- vs +/-: KWA, p = 0.008, g 1.2
<sup>c</sup> -/- vs +/+: KWA, p = 0.006, g 2.1; -/- vs +/-: KWA, p = 2E-4, g 2.6
<sup>d</sup> -/- vs +/+: 2-way ANOVA, p < 1E-9, g 1.3; -/- vs +/-: 2-way ANOVA, p < 1E-9, g 1.9

Positive current steps evoked an initial depolarizing transient in both K_V_1.8^+/+^ and K_V_1.8^−/−^ type II HCs, but the detailed time course differed (Fig. 4B). Both transient and steady-state responses were larger in K_V_1.8^−/−^, consistent with their larger R_in_ values.

Absence of K_V_1.8 increased the incidence of sharp electrical resonance in type II HCs. Electrical resonance, which manifests as ringing responses to current steps, can support receptor potential tuning (Ashmore, 1983; Fettiplace, 1987; Hudspeth and Lewis, 1988; Ramanathan and Fuchs, 2002). Larger R_in_ values made K_V_1.8^−/−^ type II HCs more prone to electrical resonance; Figure 4C.1 shows a striking example.

Median resonance quality (Q_e_, sharpness of tuning) was greater in K_V_1.8^−/−^ (1.33, n=26) than K_V_1.8^+/+^ (0.66, n=23) or K_V_1.8^+/−^ (0.59, n=44) type II HCs.

K_V_1.8 affected the time course of the initial peak in response to much larger current injections (Fig. 4D-E). Fast activation of g_A_ in control type II HCs rapidly repolarizes the membrane and then inactivates, allowing the constant input current to progressively depolarize the cell, producing a slow rebound (Fig 4D.2). This behavior has the potential to counter mechanotranduction adaptation (Vollrath and Eatock, 2003).

### K_V_1.8 immunolocalized to basolateral membranes of both type I and II HCs

If K_V_1.8 is a pore-forming subunit in the K_V_1.8-dependent conductances g_K,L_, g_A_, and g_DR_, it should localize to hair cell membranes. Figure 5 compares K_V_1.8 immunoreactivity in K_V_1.8^+/+^ and K_V_1.8^−/−^ utricles, showing specific immunoreactivity along the basolateral membranes of both hair cell types in K_V_1.8^+/+^ utricles. To identify hair cell type and localize the hair cell membrane, we used antibodies against K_V_7.4 (KCNQ4), an ion channel densely expressed in the calyceal “inner-face” membrane next to the synaptic cleft (Hurley et al., 2006; Lysakowski et al., 2011), producing a cup-like stain around type I HCs (Fig. 5A). K_V_1.8 immunoreactivity was present in hair cell membrane apposing K_V_7.4-stained calyx inner face in K_V_1.8^+/+^ utricles (Fig. 5A.1, A.2) and not in K_V_1.8^−/−^ utricles (Fig. 5A.3).

**Figure 5.**
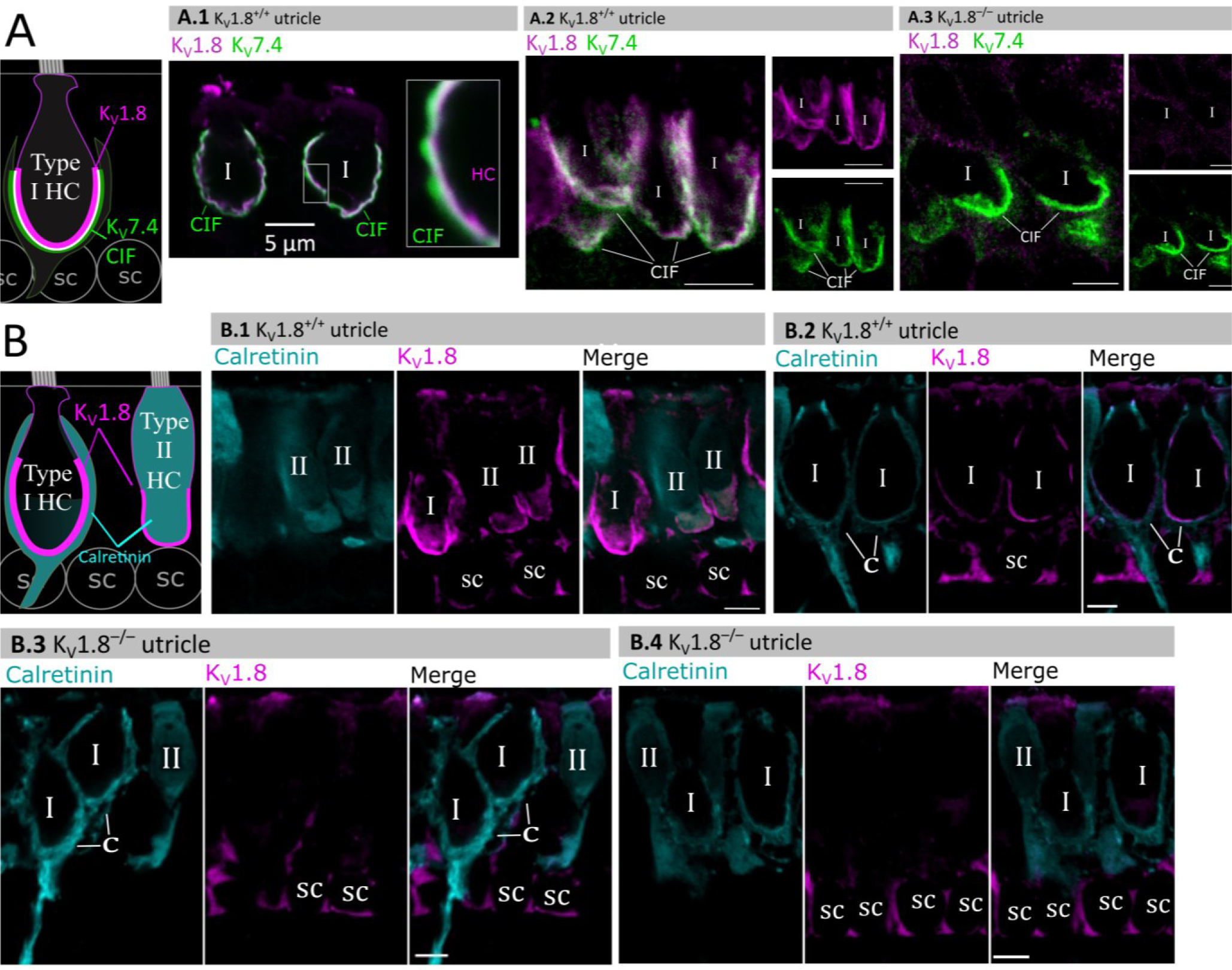
Type I and type II HC basolateral membranes show specific immunoreactivity to Kv1.8 antibody (magenta). Antibodies for K_V_7.4 (A, green) and calretinin (B, cyan) were used as counterstains for calyx membrane (Kv7.4), type II HC cytoplasm (calretinin) and cytoplasm of striolar calyx-only afferents (calretinin). (**A**) *Left,* Cartoon showing K_V_7.4 on the calyx inner face membrane (CIF) and K_V_1.8 on the type I HC membrane. SC, supporting cell nuclei. *A.1-3*, Adult mouse utricle sections. K_V_7.4 antibody labeled calyces on two K_V_1.8-positive type I HCs (**A.1**), four K_V_1.8-positive type I HCs (**A.2**), and two K_V_1.8-negative type I HCs from a K_V_1.8^−/−^ mouse (**A.3**). (**B**) *Left,* Cartoon showing cytoplasmic calretinin stain in calyx-only striolar afferents and most type II HCs, and K_V_1.8 on membranes of both HC types. In wildtype utricles, K_V_1.8 immunolocalized to basolateral membranes of type I and II HCs (**B.1**). K_V_1.8 immunolocalized to type I HCs in striola (**B.2**). Staining of supporting cell (SC) membranes by Kv1.8 antibody was non-specific, as it was present in K_V_1.8^−/−^ tissue (**B.3, B.4**).

In other experiments, we used antibodies against calretinin (Calb2), a cytosolic calcium binding protein expressed by many type II HCs and also by striolar calyx-only afferents (Desai et al., 2005; Lysakowski et al., 2011) (Fig. 5B). A hair cell is type II if it is calretinin-positive (Fig. 5B.1) or if it lacks a K_V_7.4-positive or calretinin-positive calyceal cup (Fig. 5A.2, 5B.3, rightmost cells). Hair cell identification was confirmed with established morphological indicators: for example, type II HCs tend to have basolateral processes (feet) (Pujol et al., 2014) and, in the extrastriola, more apical nuclei than type I HC.

Previously, Carlisle et al. (2012) reported K_V_1.8-like immunoreactivity in many cell types in the inner ear. In contrast, Lee et al. (2013) found that gene expression reporters indicated expression only in hair cells and some supporting cells. Here, comparison of control and null tissue showed selective expression of HC membranes, and that some supporting cell staining is non-specific.

### K_V_1.4 may also contribute to g_A_

Results with the K_V_1.8 knockout suggest that type II hair cells have an inactivating K_V_1 conductance that includes K_V_1.8 subunits. K_V_1.8, like most K_V_1 subunits, does not show fast inactivation as a heterologously expressed homomer (Lang et al., 2000; Ranjan et al., 2019; Dierich et al., 2020), nor do the K_V_1.8-dependent channels in type I HCs, as we show, and in cochlear inner hair cells (Dierich et al., 2020). K_V_1 subunits without intrinsic inactivation can produce rapidly inactivating currents by associating with K_V_β1 (KCNB1) or K_V_β3 subunits. K_V_β1 is present in type II HCs alongside K_V_β2 (McInturff et al., 2018; Jan et al., 2021; Orvis et al., 2021), which does not confer rapid inactivation (Dwenger et al., 2022).

Another possibility is that in type II HCs, K_V_1.8 subunits heteromultimerize with K_V_1.4 subunits—the only K_V_1 subunits which, when expressed as a homomer, have complete N-type (fast) inactivation (Stühmer et al., 1989). Multiple observations support this possibility. K_V_1.4 has been linked to g_A_ in pigeon type II HCs (Correia et al., 2008) and is the second-most abundant K_V_1 transcript in mammalian vestibular HCs, after K_V_1.8 (Scheffer et al., 2015). K_V_1.4 is expressed in type II HCs but not type I HCs (McInturff et al., 2018; Orvis et al., 2021), and is not found in striolar HCs (Jan et al., 2021; Orvis et al., 2021), where even in type II HCs, inactivation is slower and less extensive (Fig. 3A).

Functional heteromers form between K_V_1.4 and other K_V_1.x and/or K_V_β1 (Imbrici et al., 2006; Correia et al., 2008; Al-Sabi et al., 2011). Although K_V_1.4 and K_V_1.8 heteromers have not been studied directly, g_A_’s inactivation time course (τ_Fast,Inac_ of ∼30 ms +30 mV, Fig. 3A) and voltage dependence (V_half_ –41 mV, Fig. 6B) are consistent with these other K_V_1.4-containing heteromers.

**Figure 6.**
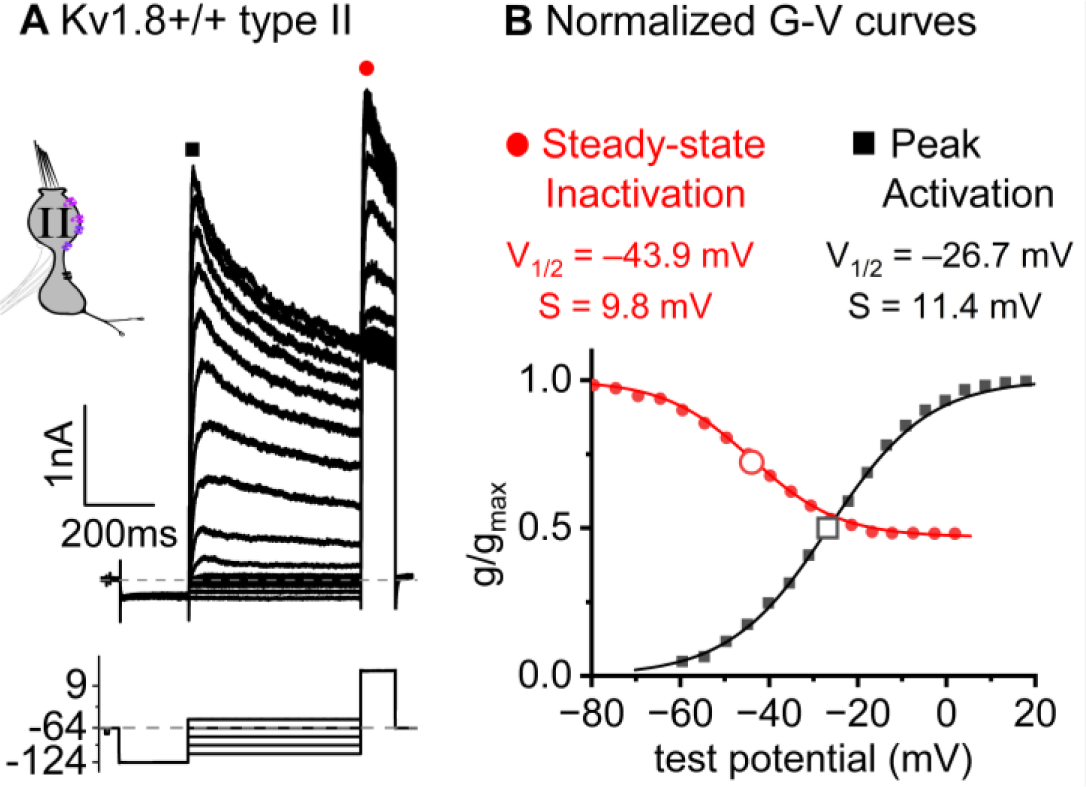
Inactivation curve of g_A_ in extrastriolar type II HCs. (**A**) Modified voltage protocol measured accumulated steady-state inactivation in the tail potential. 100 μM ZD7288 in bath prevented contamination by HCN current. (**B**) Voltage dependence of g_A_’s steady-state inactivation (h_∞_ curve) and peak activation are consistent with K_V_1.4 heteromers. H_∞_ curves have overlapped normalized activation (peak conductance, black squares) and inactivation (tail conductance, red circles) G-V data points. *Curves,* Boltzmann fits (Eq. 1). *Average fit parameters* from K_V_1.8^+/+,+/−^ type II HCs, P40-P210, median P94. Inactivation: V_half_, –42 ±2 mV (n=11); S, 11 ± 1 mV. Activation: V_half_, –23 ± 1 mV (n=11); S, 11.2 ± 0.4 mV.

### K_V_7 channels contribute a small delayed rectifier in type I and type II hair cells

In K_V_1.8^−/−^ HCs, absence of I_K,L_ and I_A_ revealed smaller delayed rectifier K^+^ currents that, unlike I_K,L_, activated positive to resting potential and, unlike I_A_, lacked fast inactivation. Candidate channels include members of the K_V_7 (KCNQ, M-current) family, which have been identified previously in rodent vestibular HCs (Kharkovets et al., 2000; Rennie et al., 2001; Hurley et al., 2006; Scheffer et al., 2015).

We test for K_V_7 contributions in K_V_1.8^−/−^ type I HCs, K_V_1.8^−/−^ type II HCs, and K_V_1.8^+/+,+/−^ type II HCs of multiple ages by applying XE991 at 10 µM (Fig. 7A), a dose selective for K_V_7 channels (Brown et al., 2002) and close to the IC_50_ (Alexander et al., 2019). In K_V_1.8^−/−^ HCs of both types, 10 µM XE991 blocked about half of the residual K_V_ conductance (Fig. 7B.1), consistent with K_V_7 channels conducting most or all of the non-K_V_1.8 delayed rectifier current. In all tested HCs (P8-355, median P224), the XE991-sensitive conductance did not inactivate substantially within 200 ms at any voltage, consistent with K_V_7.2, 7.3, 7.4, and 7.5 currents (Wang, 1998; Kubisch et al., 1999; Schroeder et al., 2000; Jensen et al., 2007; Xu et al., 2007). We refer to this component as g_DR_(K_v_7). The voltage dependence and Gmax density (G_max_/C_m_) of g_DR_(K_v_7) were comparable across HC types and genotypes (Figure 7B.2-4). Although K_V_7.4 was not detectable in HCs during immunostaining (Fig. 5), K_V_7.4 has been shown in type I HCs with immunogold labeling (Kharkovets et al., 2000; Hurley et al., 2006).

**Figure 7.**
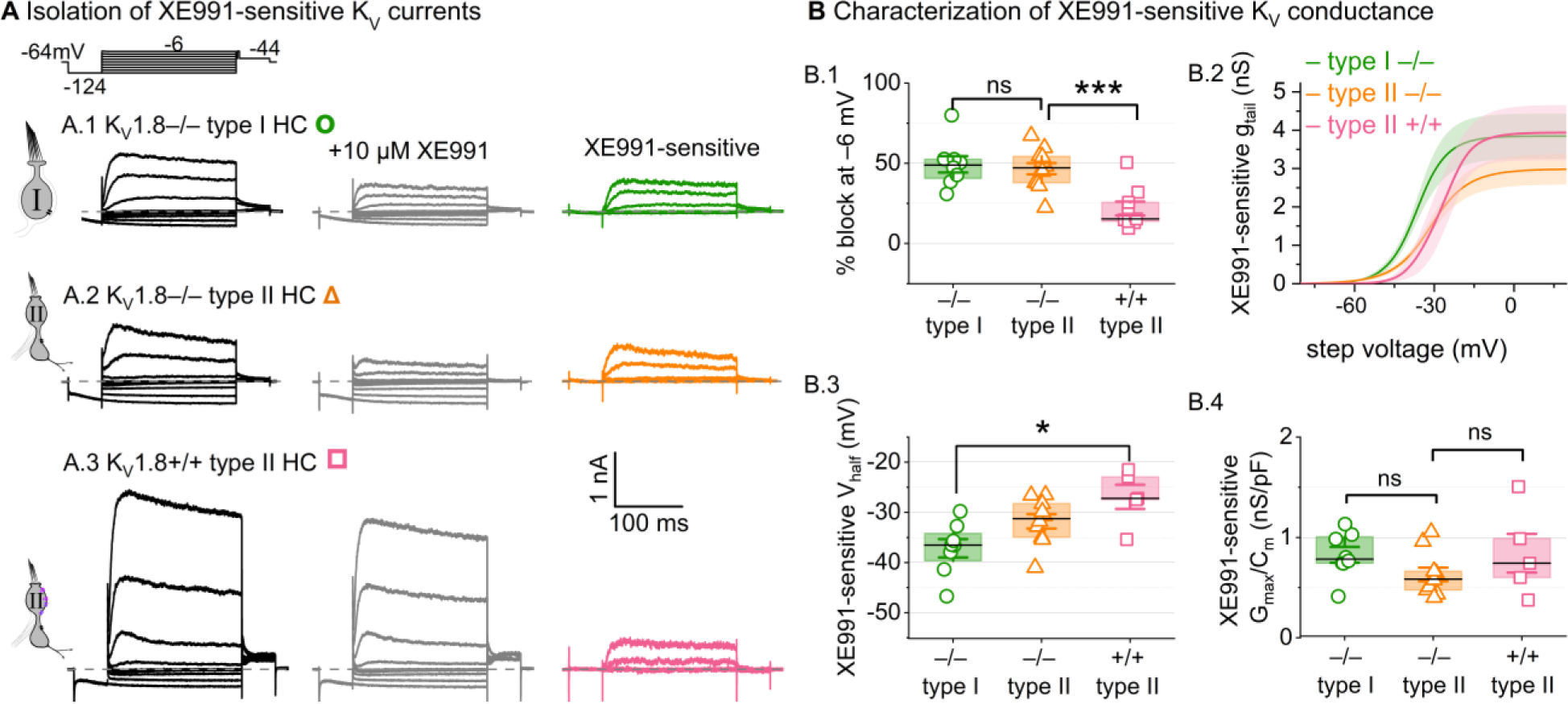
A K_V_7-selective blocker, XE991, reduced residual delayed rectifier currents in K_V_1.8^−/−^ type I and II HCs. **(A)** XE991 (10 μM) partly blocked similar delayed rectifier currents in type I and type II K_V_1.8^−/−^ HCs and a type II K_V_1.8^+/+^ HC. **(B)** Properties of XE991-sensitive conductance, g_DR_(K_V_7). (**B.1**) % Block of steady-state current. (**B.2**) tail G-V curves for K_V_1.8^−/−^ type I HCs (n=8), K_V_1.8^−/−^ type II HCs (9), and K_V_1.8^+/+^ type II HCs (5); mean ± SEM. (**B.3**) V_half_ was less negative in K_V_1.8^+/+^ type II than K_V_1.8^−/−^ type I HC (p = 0.01, KWA). (**B.4**) Conductance density was similar in all groups (ANOVA, non-significant at 40% power (*left*), 20% power (*right*).

These results are consistent with similar K_V_7 channels contributing a relatively small delayed rectifier in both HC types. In addition, the similarity of XE991-sensitive currents of K_V_1.8^+/+^ and K_V_1.8^−/−^ type II HCs indicates that knocking out K_V_1.8 did not cause general effects on ion channel expression. We did not test XE991 on K_V_1.8^+/+,+/−^ type I HCs because g_K,L_ runs down in ruptured patch recordings (Rüsch and Eatock, 1996a; Chen and Eatock, 2000; Hurley et al., 2006), which could contaminate the XE991-sensitive conductance obtained by subtraction.

In one striolar K_V_1.8^−/−^ type I HC, XE991 also blocked a small conductance that activated negative to rest (Suppl. Fig. 5A-B). This conductance (V_half_ ∼= –100 mV, Suppl. Fig. 5C) was detected only in K_V_1.8^−/−^ type I HCs from the striola (5/23 vs. 0/45 extrastriolar). The V_half_ and τ_deactivation_ were similar to values reported for K_V_7.4 channels in cochlear HCs (Wong et al., 2004; Dierich et al., 2020). This very negatively activating K_V_7 conductance coexisted with the larger more positively activating K_V_7 conductance (Suppl. Fig. 5C) and was too small (<0.5 nS/pF) to contribute significantly to g_K,L_ (∼10-100 nS/pF, Fig. 1D).

### Other channels

While our data are consistent with K_V_1.8- and K_V_7-containing channels carrying most of the outward-rectifying current in mouse utricular hair cells, there is evidence in other preparations for additional channels, including K_V_11 (KCNH, Erg) channels in rat utricular type I hair cells (Hurley et al., 2006) and BK (KCNMA1) channels in rat utricle and rat and turtle semicircular canal hair cells (Schweizer et al., 2009; Contini et al., 2020).

BK is expressed in mouse utricular hair cells (McInturff et al., 2018; Jan et al., 2021; Orvis et al., 2021). However, Ca^2+^-dependent currents have not been observed in mouse utricular HCs, and we found little to no effect of the BK-channel blocker iberiotoxin at a dose (100 nM) well beyond the IC_50_: percent blocked at –30 mV was 2 ± 6% (3 K_V_1.8^−/−^ type I HCs); 1 ± 5% (5 K_V_1.8^+/+,+/−^ type II HCs); 7% and 14% (2 K_V_1.8^−/−^ type II HCs). We also did not see N-shaped I-V curves typical of many Ca^2+^-dependent K^+^ currents. In our ruptured-patch recordings, Ca^2+^-dependent BK currents and erg channels may have been eliminated by wash-out of the hair cells’ small Ca_V_ currents (Bao et al., 2003) or cytoplasmic second messengers (Hurley et al., 2006).

To check whether the constitutive K_V_1.8 knockout has strong non-specific effects on channel trafficking, we examined the summed HCN and fast inward rectifier currents (I_H_ and I_Kir_) at –124 mV, and found them similar across genotypes (Suppl. Fig. 6). The g_K,L_ knockout allowed identification of zonal differences in I_H_ and I_Kir_ in type I HCs, previously examined in type II HCs (Masetto and Correia, 1997; Levin and Holt, 2012). In type I HCs from both control and null utricles, I_H_ and I_Kir_ were less prevalent in striola than extrastriola, and, when present, the combined inward current was smaller (Suppl. Fig. 6B).

## Discussion

We have shown that constitutive knockout of K_V_1.8 eliminated g_K,L_ in type I HCs, and g_A_ and much of g_DR_ in type II HCs. K_V_1.8 immunolocalized specifically to the basolateral membranes of type I and II HCs. We conclude that K_V_1.8 is a pore-forming subunit of g_K,L_, g_A_, and part of g_DR_ [g_DR_(K_V_1.8)]. We suggest that fast inactivation of g_A_ may arise from heteromultimerization of non-inactivating K_V_1.8 subunits and inactivating K_V_1.4 subunits. Finally, we showed that a substantial component of the residual delayed rectifier current in both type I and type II HCs comprises K_V_7 channels.

K_V_1.8 is expressed in hair cells from mammalian cochlea (Dierich et al., 2020), avian utricle (Scheibinger et al., 2022), and zebrafish (Erickson and Nicolson, 2015). Our work suggests that in anamniotes, which lack type I cells and g_K,L_, K_V_1.8 contributes to g_A_ and g_DR_, which are widespread in vertebrate HCs (reviewed in <u>Meredith and Rennie, 2016)</u>. K_V_1.8 expression has not been detected in rodent brain but is reported in the pacemaker nucleus of weakly electric fish (Smith et al., 2018).

### K_V_1.8 subunits may form homomultimers to produce g_K,L_ in type I hair cells

Recent single-cell expression studies on mouse utricles (McInturff et al., 2018; Jan et al., 2021; Orvis et al., 2021) have detected just one K_V_1 subunit, K_V_1.8, in mouse type I HCs. Given that K_V_1.8 can only form multimers with K_V_1 family members, and given that g_K,L_ channels are present at very high density (∼150 per μm^2^ in rat type I, Chen and Eatock, 2000), it stands to reason that most or all of the channels are K_V_1.8 homomers. Other evidence is consistent with this proposal. g_K,L_ (Rüsch and Eatock, 1996a) and heterologously expressed K_V_1.8 homomers in oocytes (Lang et al., 2000) are non-inactivating and blocked by millimolar Ba^2+^ and 4-aminopyridine and >10 mM TEA. Unlike channels with K_V_1.1, K_V_1.2, and K_V_1.6 subunits, g_K,L_ is not sensitive to 10 nM α-dendrotoxin (Rüsch and Eatock, 1996a). g_K,L_ and heterologously expressed K_V_1.8 channels have similar single-channel conductances (∼20 pS for g_K,L_ at positive potentials, Chen and Eatock, 2000; 11 pS in oocytes, Lang et al., 2000). g_K,L_ is inhibited—or positively voltage-shifted— by cGMP (Behrend et al., 1997; Chen and Eatock, 2000), presumably via the C-terminal cyclic nucleotide binding domain of K_V_1.8.

A major novel property of g_K,L_ is that it activates 30-60 mV negative to type II K_V_1.8 conductances and most other low-voltage-activated K_V_ channels (Ranjan et al., 2019). The very negative activation range is a striking difference between g_K,L_ and known homomeric K_V_1.8 channels. Heterologously expressed homomeric K_V_1.8 channels have an activation V_half_ of –10 to 0 mV (*X. laevis* oocytes, <u>Lang et al., 2000</u>; CHO cells, Dierich et al., 2020). In cochlear inner HCs, currents attributed to K_V_1.8 (by subtraction of other candidates) have a near-zero activation V_half_ (–4 mV, Dierich et al., 2020).

Possible factors in the unusually negative voltage dependence of g_K,L_ include:

1. *elevation of extracellular K^+^* by the enveloping calyceal terminal, unique to type I HCs (Lim et al., 2011; Contini et al., 2012; Spaiardi et al., 2020; Govindaraju et al., 2023). High K^+^ increases conductance though g_K,L_ channels (Contini et al., 2020), perhaps through K^+^-mediated relief of C-type inactivation (López-Barneo et al., 1993; Baukrowitz and Yellen, 1995). We note, however, that g_K,L_ is open at rest even in neonatal mouse utricles cultured without innervation (Rüsch et al., 1998) and persists in dissociated type I HCs (Chen and Eatock, 2000; Hurley et al., 2006).
2. *The high density of g_K,L_* (∼50 nS/pF in striolar K_V_1.8^+/+^ HCs) implies close packing of channels, possibly represented by particles (12-14 nm) seen in freeze-fracture electron microscopy of the type I HC membrane (Gulley and Bagger-Sjöbäck, 1979; Sousa et al., 2009). Such close channel packing might hyperpolarize *in situ* voltage dependence of g_K,L_, as proposed for K_V_7.4 channels in outer hair cells (Perez-Flores et al., 2020). Type I HC-specific partners that may facilitate this close packing include ADAM11 (McInturff et al., 2018), which clusters presynaptic K_V_1.1 and K_V_1.2 to enable ephaptic coupling at a cerebellar synapse (Kole et al., 2015).
3. *Modulation by accessory subunits*. Type I HCs express K_V_β1 (McInturff et al., 2018; Orvis et al., 2021), an accessory subunit that can confer fast inactivation and hyperpolarize activation V_half_ by ∼10 mV. K_V_β1 might interact with K_V_1.8 to shift voltage dependence negatively. Arguments against this possibility include that g_K,L_ lacks fast inactivation (Rüsch and Eatock, 1996a; Hurley et al., 2006; Spaiardi et al., 2017) and that cochlear inner hair cells co-express K_V_1.8 and K_V_β1 (Liu et al., 2018) but their K_V_1.8 conductance has a near-0 V_half_ (Dierich et al., 2020).

### K_V_1.8 subunits may combine with different subunits to produce g_A_ and K_V_1.8-dependent g_DR_ in type II HCs

The K_V_1.8-dependent conductances of type II HCs vary in their fast and slow inactivation. In not showing fast inactivation (Lang et al., 2000; Ranjan et al., 2019; Dierich et al., 2020), heterologously expressed K_V_1.8 subunits resemble most other K_V_1 family subunits, with the exception of K_V_1.4 (for comprehensive review, see Ranjan et al., 2019). K_V_1.4 is a good candidate to provide fast inactivation based on immunolocalization and voltage dependence (Figs. 4, 6). We suggest that g_A_ and g_DR_(K_V_1.8) are K_V_1.8-containing channels that may include a variable number of K_V_1.4 subunits and K_V_β2 and K_V_β1 accessory subunits.

K_V_1.4-K_V_1.8 heteromeric assembly could account for several related observations. The faster τ_Inact,Fast_ in K_V_1.8^+/−^ relative to K_V_1.8^+/+^ type II HCs (Fig. 3A.3, Suppl. Fig. 2A.1) could reflect an increased ratio of K_V_1.4 to K_V_1.8 subunits and therefore more N-terminal inactivation domains per heteromeric channel. Zonal variation in the extent and speed of N-type inactivation (Fig. 3A) might arise from differential expression of K_V_1.4. The small fast-inactivating conductance in ∼20% of extrastriolar K_V_1.8^−/−^ type II HCs (Suppl. Fig. 4) might flow through K_V_1.4 homomers.

Fast inactivation may also receive contribution from K_V_β subunits. K_V_β1 is expressed in type II HCs (McInturff et al., 2018; Jan et al., 2021; Orvis et al., 2021), and, together with K_V_1.4, has been linked to g_A_ in pigeon vestibular HCs (Correia et al., 2008). K_V_β2, also expressed in type II HCs (McInturff et al., 2018; Orvis et al., 2021), does not confer fast inactivation.

We speculate that g_A_ and g_DR_(K_V_1.8) have different subunit composition: g_A_ may include heteromers of K_V_1.8 with other subunits that confer rapid inactivation, while g_DR_(K_V_1.8) may comprise homomeric K_V_1.8 channels, given that they do not have N-type inactivation.

### K_V_1.8 relevance for vestibular function

In both type I and type II utricular HCs, K_V_1.8-dependent channels strongly shape receptor potentials in ways that promote temporal fidelity rather than electrical tuning (Lewis, 1988), consistent with the utricle’s role in driving reflexes that compensate for head motions as they occur. This effect is especially pronounced for type I HCs, where the current-step evoked voltage response reproduces the input with great speed and linearity (Fig. 2).

g_K,L_ dominates passive membrane properties in mature K_V_1.8^+/+,+/−^ type I HCs such that K_V_1.8^−/−^ type I HCs are expected to have receptor potentials with higher amplitudes but lower low-pass corner frequencies, closer to those of type II HCs and immature HCs of all types (Correia et al., 1996; Rüsch and Eatock, 1996a; Songer and Eatock, 2013). In K_V_1.8^−/−^ epithelia, we expect the lack of a large basolateral conductance open at rest to reduce the speed and gain of non-quantal transmission, which depends on K^+^ ion efflux from the type I HC to change electrical and K^+^ potentials in the synaptic cleft (Govindaraju et al., 2023). In hair cells, K^+^ enters the mechanosensitive channels of the hair bundle from the K^+^-rich apical endolymph and exits through basolateral potassium conductances into the more conventional low-K+ perilymph. For the type I-calyx synapse, having in the hair cell a large, non-inactivating K^+^ conductance open across the physiological range of potentials avoids channel gating time and allows for instantaneous changes in current into the cleft and fast afferent signaling (Pastras et al., 2023).

In contrast, mature type II HCs face smaller synaptic contacts and have K_V_1.8-dependent currents that are not substantially activated at resting potential. They do affect the time course and gain of type II HC responses to input currents, speeding up depolarizing transients, producing a repolarizing rebound during the step, and reducing resonance.

Type I and II vestibular hair cells are closely related, such that adult type II HCs acquire type I-like properties upon deletion of the transcription factor Sox2 (Stone et al., 2021). In normal development of the two cell types, the Kcna10 gene generates biophysically distinct and functionally different ion channels, presenting a natural experiment in functional differentiation of sensory receptor cells.

## Materials and methods

### Preparation

All procedures for handling animals followed the NIH Guide for the Care and Use of Laboratory Animals and were approved by the Institutional Animal Care and Use Committees of the University of Chicago and the University of Illinois Chicago. Most mice belonged to a transgenic line with a knockout allele of *Kcna10* (referred to here as K_V_1.8^−/−^). Our breeding colony was established with a generous gift of such animals from Sherry M. Jones and Thomas Friedman. These animals are described in their paper (Lee et al., 2013). Briefly, the Texas A&M Institute for Genomic Medicine generated the line on a C57BL/6;129SvEv mixed background by replacing Exon 3 of the *Kcna10* gene with an IRES-bGeo/Purocassette. Mice in our colony were raised on a 12:12h light-dark cycle with access to food and water *ad libitum*.

Semi-intact utricles were prepared from ∼150 male and ∼120 female mice, postnatal days (P) 5-375, for same-day recording. Hair cell K_V_ channel data were pooled across sexes as most results did not appear to differ by sex; an exception was that g_K,L_ had a more negative V_half_ in males (Suppl. Table 1), an effect not clearly related to age, copy number, or other properties of the activation curve.

Preparation, stimulation, and recording methods followed our previously described methods for the mouse utricle (Vollrath and Eatock, 2003). Mice were anesthetized through isoflurane inhalation. After decapitation, each hemisphere was bathed in ice-cold, oxygenated Liebowitz-15 (L15) media. The temporal bone was removed, the labyrinth was cut to isolate the utricle, and the nerve was cut close to the utricle. The utricle was treated with proteinase XXIV (100 μg/mL, ∼10 mins, 22°C) to facilitate removal of the otoconia and attached gel layer and mounted beneath two glass rods affixed at one end to a coverslip.

### Electrophysiology

We used the HEKA Multiclamp EPC10 with Patchmaster acquisition software, filtered by the integrated HEKA filters: a 6-pole Bessel filter at 10 kHz and a second 4-pole Bessel filter at 5 kHz, and sampled at 10-100 kHz. Recording electrodes were pulled (PC-100, Narishige) from soda lime glass (King’s Precision Glass R-6) wrapped in paraffin to reduce pipette capacitance. Internal solution contained (in mM) 135 KCl, 0.5 MgCl_2_, 3 MgATP, 5 HEPES, 5 EGTA, 0.1 CaCl_2_, 0.1 Na-cAMP, 0.1 LiGTP, 5 Na_2_CreatinePO_4_ adjusted to pH 7.25 and ∼280 mmol/kg by adding ∼30 mM KOH. External solution was Liebowitz-15 media supplemented with 10 mM HEPES (pH 7.40, 310 ± 10 mmol/kg). Recording temperature was 22-25°C. Pipette capacitance and membrane capacitance transients were subtracted during recordings with Patchmaster software. Series resistance (8-12 MΩ) was measured after rupture and compensated 60-80% with the amplifier, to final values of ∼2 MΩ. Potentials are corrected for remaining (uncompensated) series resistance and liquid junction potential of ∼+4 mV, calculated with LJPCalc software (Marino et al., 2014).

K_V_1.8^−/−^ hair cells appeared healthy in that cells had resting potentials negative to –50 mV, cells lasted a long time (20-30 minutes) in ruptured patch recordings, membranes were not fragile, and extensive blebbing was not seen. Type I HCs with g_K,L_ were transiently hyperpolarized to –90 mV to close g_K,L_ enough to increase R_input_ above 100 MΩ, as needed to estimate series resistance and cell capacitance. The average resting potential, V_rest_, was –87 mV ± 1 (41), similar to the calculated E_K_ of –86.1 mV, which is not surprising given the large K^+^ conductance of these cells. V_rest_ is likely more positive *in vivo*, where lower endolymphatic Ca^2+^ increases standing inward current through MET channels.

Voltage protocols to characterize K_V_ currents differed slightly for type I and II HCs. In standard protocols, the cell is held at a voltage near resting potential (–74 mV in type I and –64 mV in type II), then jumped to –124 mV for 200 ms in type I HCs in order to fully deactivate g_K,L_ and 50 ms in type II HCs in order to remove baseline inactivation of g_A_. The subsequent iterated step depolarizations lasted 500 ms in type I HCs because g_K,L_ activates slowly (Wong et al., 2004) and 200 ms in type II HCs, where K_V_ conductances activate faster. The 50-ms tail voltage was near the reversal potential of HCN channels (–44 mV in mouse utricular hair cells, Rüsch et al., 1998) to avoid HCN current contamination.

G-V (activation) parameters for control type I cells may be expected to vary across experiments on semi-intact (as here), organotypically cultured and denervated (Rüsch et al.,1998), or dissociated-cell preparations, reflecting variation in retention of the calyx (Discussion) and voltage step durations (Wong et al., 2004) which elevate K^+^ concentration around the hair cell. Nevertheless, the values we obtained for type I and type II HCs resemble values recorded elsewhere, including experiments in which extra care was taken to avoid extracellular K^+^ accumulation (Spaiardi et al., 2017, 2020). The effects of K^+^ accumulation on g_K,L_’s steady-state activation curves are small because the operating range is centered on E_K_ and can be characterized with relatively small currents (Fig. 1A).

### Pharmacology

Drug-containing solutions were locally with BASI Bee Hive syringes at a final flow rate of 20 μL/min and a dead time of ∼30 s. Global bath perfusion was paused during drug perfusion and recording, and only one cell was used per utricle. Aliquots of test agents in solution were prepared, stored at –20°C, and thawed and added to external solution on the recording day (see Key Resources Table).

**Table 6.** Key Resources Table.

| Antibody | Source | Catalog Number | Lot Number | Dilution |
| --- | --- | --- | --- | --- |
| Rabbit anti-K<sub>v</sub>1.8 | Alomone | APC-157 | 0102 | 1:200 or 1:400 |
| Mouse anti-K<sub>v</sub>1.4 | NeuroMab | P15385 | 5HK-05 | 1:400 |
| Mouse IgG2a-conjugated anti-Tuj1 | Covance | MMS-435P | B205808 | 1:300 |
| Goat anti-calretinin | Millipore | AB1550 | 9669 | 1:600 |
| Mouse IgG1-conjugated anti-K<sub>v</sub>7.4 | NeuroMab | 2HK-65 |  | 1:200 |
| Donkey anti-Rabbit Secondary Antibody, Alexa Fluor 594 | Invitrogen | A21207 | 8652 | 1:200 |
| Donkey anti-Goat 488 nm | Invitrogen | A21125 | 1920483 | 1:200 |
| Goat anti-Mouse IgG1 594 | Invitrogen | A11055 | 1869589 | 1:200 |
| Chemicals and Peptides | Source | Catalog Number | Diluent |  |
| Iberiotoxin | Alomone | STI-400 | Water |  |
| XE991 dihydrochloride | Sigma | X2254 | Water |  |
| ZD7288 | Tocris | APN18035-2 | Water |  |
| Bovine serum albumin | Fisher | BP671 | water |  |

### Analysis

Data analysis was performed with software from OriginLab (Northampton, MA) and custom MATLAB scripts using MATLAB fitting algorithms.

### Fitting voltage dependence and time course of conductances

*G-V curves.* Current was converted to conductance (G) by dividing by driving force (V – E_K_; E_K_ calculated from solutions). For type I HCs, tail G-V curves were generated from current 1 ms after the end of the iterated voltage test step. For type II HCs, peak G-V curves were generated from peak current during the step and steady-state G-V curves were generated from current 1 ms before the end of a 200 ms step. Sigmoidal voltage dependence of G-V curves was fit with the first-order Boltzmann equation (Eq. 1).

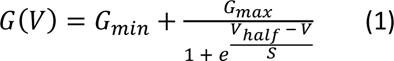

V_half_ is the midpoint and S is the slope factor, inversely related to curve steepness near activation threshold.

*Activation time course of type II HCs.* For type II HCs lacking fast inactivation, outward current activation was fit with Eq. 2.

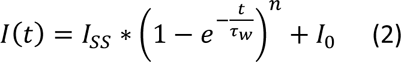

I_SS_ is steady-state current, τ_w_ is activation time constant, n is the state factor related to the number of closed states (typically constrained to 3), and I_o_ is baseline current.

To measure activation and inactivation time course of g_A_, we used Eq. 3 to fit outward K^+^ currents evoked by steps from –125 mV to above –50 mV (Rothman and Manis, 2003b).

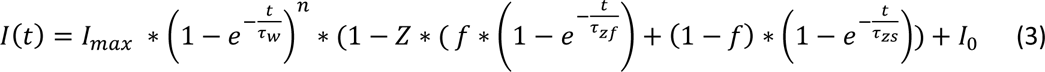

Z is total steady-state inactivation (0 ≤ Z < 1 means incomplete inactivation, which allows the equation to fit non-inactivating delayed rectifier currents), f is the fraction of fast inactivation relative to total inactivation, I_max_ is maximal current, τ_zf_ and τ_zs_ are the fast and slow inactivation time constants. We chose to compare fit parameters at 30 ± 2 mV (91), where fast and slow inactivation were consistently separable and g_A_ was maximized. In most K_V_1.8^−/−^ and some striolar K_V_1.8^+/+,+/−^ cells, where fast inactivation was absent and adjusted R^2^ did not improve on a single-exponential fit by >0.01, we constrained f in Eq. 3 to 0 to avoid overfitting.

For *Peak* G-V relations, peak conductance was taken from fitted curves (Eqs. 2 and 3). To construct ‘*Steady-state’* G-V relations, we used current at 200 ms (6 ± 1 % (94) greater than steady-state estimated from fits to Eq. 3 (Fig. 3C-D)).

Percent inactivation was calculated at 30 mV with Eq. 4 :

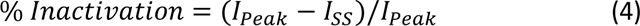

I_Peak_ is maximal current, and I_SS_ is current at the end of a 200 ms voltage step.

The electrical resonance of type II HCs was quantified by fitting voltage responses to current injection steps (Songer and Eatock, 2013). We fit Eq. 5, a damped sinusoid, to the voltage trace from half-maximum of the initial depolarizing peak until the end of the current step.

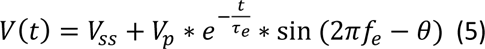

V_SS_ is steady-state voltage, V_p_ is the voltage of the peak response, τ_e_ is the decay time constant, f_e_ is the fundamental frequency, and θ is the phase angle shift.

Quality factor, Q_e_, was calculated with Eq. 6 (Crawford and Fettiplace, 1981).

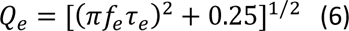

### Statistics

We give means ± SEM for normally-distributed data, and otherwise, median and range. Data normality was assessed with the Shapiro-Wilk test for n<50 and the Kolmogorov-Smirnov test for n>50. To assess homogeneity of variance we used Levene’s test. With homogeneous variance, we used two-way ANOVA for genotype and zone with the posthoc Tukey’s test. When variance was non-homogeneous, we used one-way Welch ANOVA with the posthoc Games-Howell test. For data that were not normally distributed, we used the non-parametric one-way Kruskal-Wallis ANOVA (KWA) with posthoc Dunn’s test. Effect size is Hedge’s g (g). For age dependence, we used partial correlation coefficients controlling for genotype and zone. Statistical groups may have different median ages, but all have overlapping age ranges. In figures, asterisks represent p-value ranges as follows: *, p<0.05; **, p<0.01; ***, p<0.001; ****, p<0.0001.

### Immunohistochemistry

Mice were anesthetized with Nembutal (80 mg/kg), then perfused transcardially with 40mL of physiological saline containing heparin (400 IU), followed by 2 mL/g body weight fixative (4% paraformaldehyde, 1% picric acid, and 5% sucrose in 0.1 M phosphate buffer at pH 7.4, sometimes with 1% acrolein). Vestibular epithelia were dissected in phosphate buffer, and tissues were cryoprotected in 30% sucrose-phosphate buffer overnight at 4°C. Otoconia were dissolved with Cal-Ex (Fisher Scientific) for 10 min. Frozen sections (35 μm) were cut with a sliding microtome. Immunohistochemistry was performed on free-floating sections. Tissues were first permeabilized with 4% Triton X-100 in PBS for 1 h at room temperature, then incubated with 0.5% Triton X-100 in a blocking solution of 0.5% fish gelatin and 1% BSA for 1 h at room temperature. Sections were incubated with 2-3 primary antibodies for 72 h at 4°C and with 2-3 secondary antibodies. Sections were rinsed with PBS between and after incubations and mounted on slides in Mowiol (Calbiochem).

## Abbreviations

g_A_: A-type (inactivating) K_V_ conductance in type II HCs
g_DR_: delayed rectifier K^+^ conductance
g_K,L_: low-voltage-activated K^+^ conductance in type I
HCs HC: hair cell
K_V_: voltage-gated K^+^ conductance

## Data Availability

Data used in this study are available on Dryad (https://doi.org/10.5061/dryad.37pvmcvrw).

## Acknowledgements

This study was supported by NIH grant R01 DC012347 to RAE and AL and an NSF Graduate Research Fellowship to HRM. We thank Drs. Thomas Friedman and Sherri Jones for the generous gift of the K_V_1.8^−/−^ mouse line, and Drs. Zheng-Yi Chen and Deborah I. Scheffer for bringing the expression of this subunit in mouse vestibular hair cells to our attention.

We acknowledge Dr. Vicente Lumbreras for insights from his prior experiments on g_A_ in mouse utricular hair cells, and thank him for helpful further discussions.

We acknowledge Steven D. Price for his help with immunocytochemistry.

We thank Drs. Rebecca Lim and Ebenezer Yamoah for their critical feedback on the manuscript, and Drs. Rob Raphael and Aravind Chenrayan Govindaraju for feedback and many helpful discussions. We thank Drs. Joe Burns, Gabi Pregernig, and Lars Becker (Decibel Therapeutics, Inc.) for helpful discussions.

## Author Contributions

HRM designed and performed experiments, analyzed data, and wrote the paper; RAE helped design experiments, analyze data, and write the paper; AL performed immunohistochemistry experiments and reviewed and edited the manuscript.

## Supplemental Figures

**Supplemental Figure 1.**
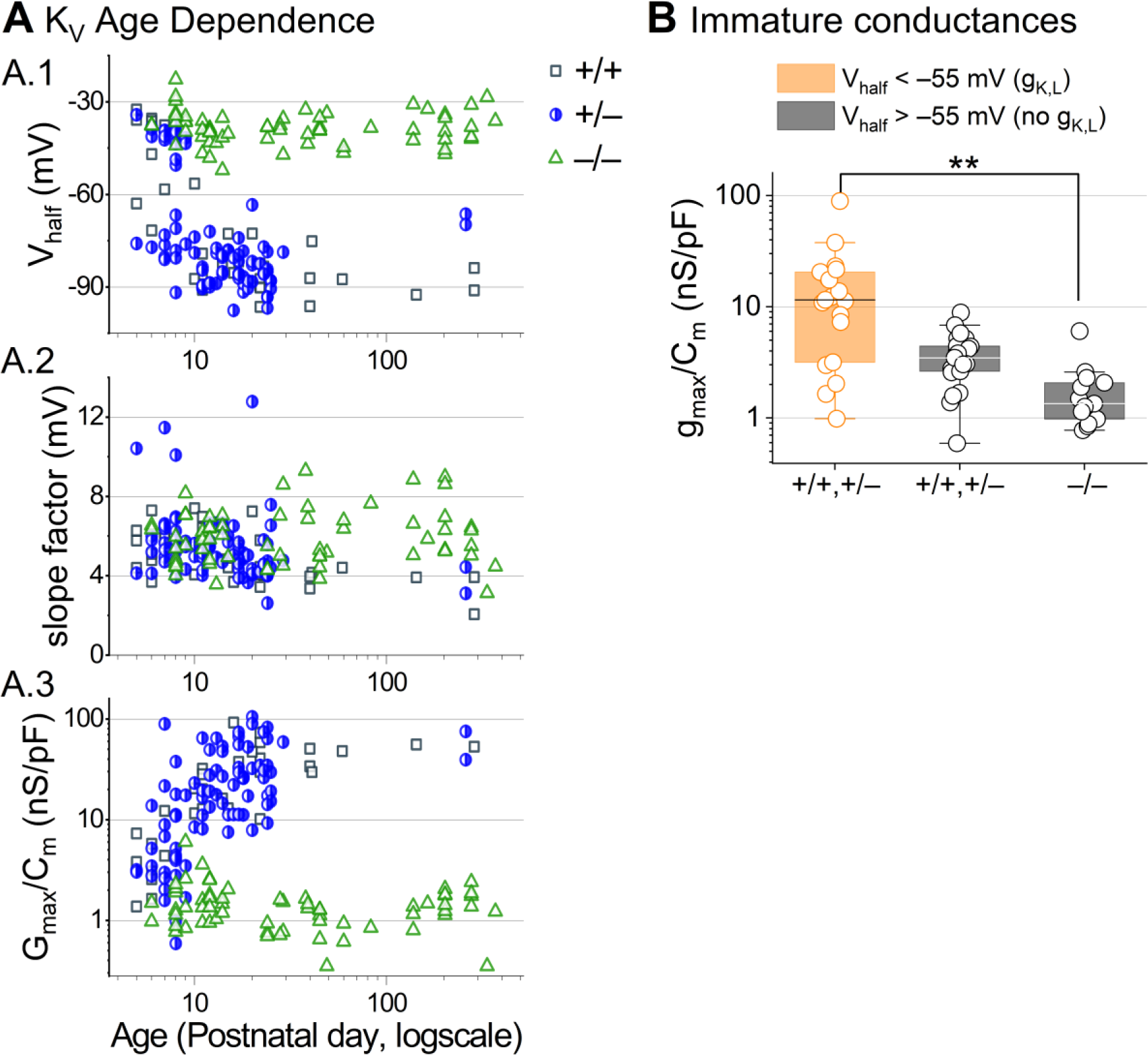
Developmental changes in type I HC K_V_ conductances. (**A**) Parameters from Boltzmann fits of tail G-V relations for type I HCs plotted against age. (**B**) Conductance density is similar in young (P5-P10) type I HCs that lack g_K,L_. g_K,L_ is defined here as having a V_half_ negative to −55 mV . K_V_1.8^+/+,+/−^ *with* g_K,L_, 17 ± 5 nS/pF (19); K_V_1.8^+/+,+/−^ *without* g_K,L_, 3.7 ± 0.4 nS/pF (22); K_V_1.8^−/−^, 1.8 ± 0.4 nS/pF (13). K_V_1.8^+/+,+/−^ *with* g_K,L_ vs. K_V_1.8^−/−^: p = 0.007, KWA, g 1.0.

**Supplemental Figure 2.**
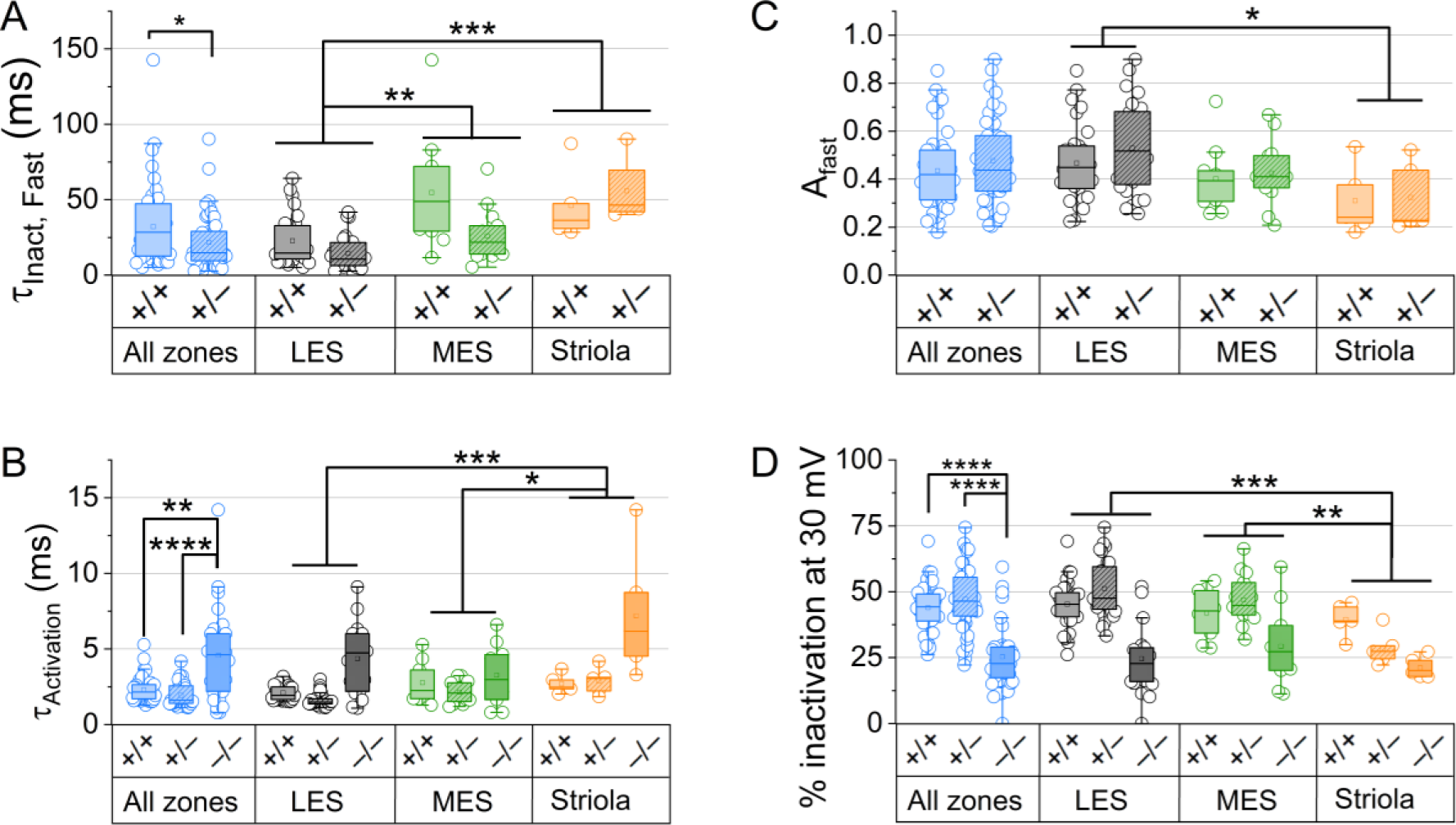
For type II HCs older than P12, K_V_ conductance activation and inactivation differed across zones and genotypes. (A) τ_inact,Fast_ at 30 mV was fastest in LES in K_V_1.8^+/+^ and K_V_1.8^+/−^ HCs, and faster in K_V_1.8^+/−^ than K_V_1.8^+/+^ HCs (see Table 3 for p-values). (B) Fast inactivation was a larger fraction of the total in LES than striola. (C) τ_Act_ at 30 mV was slower in K_V_1.8^−/−^ than K_V_1.8^+/+^ and K_V_1.8^+/−^, and slower in striola than LES and MES. (D) Percent inactivation at 30 mV was lowest in striola (zone effect), and lowest in K_V_1.8^−/−^ HCs (genotype effect).

**Supplemental Figure 3.**
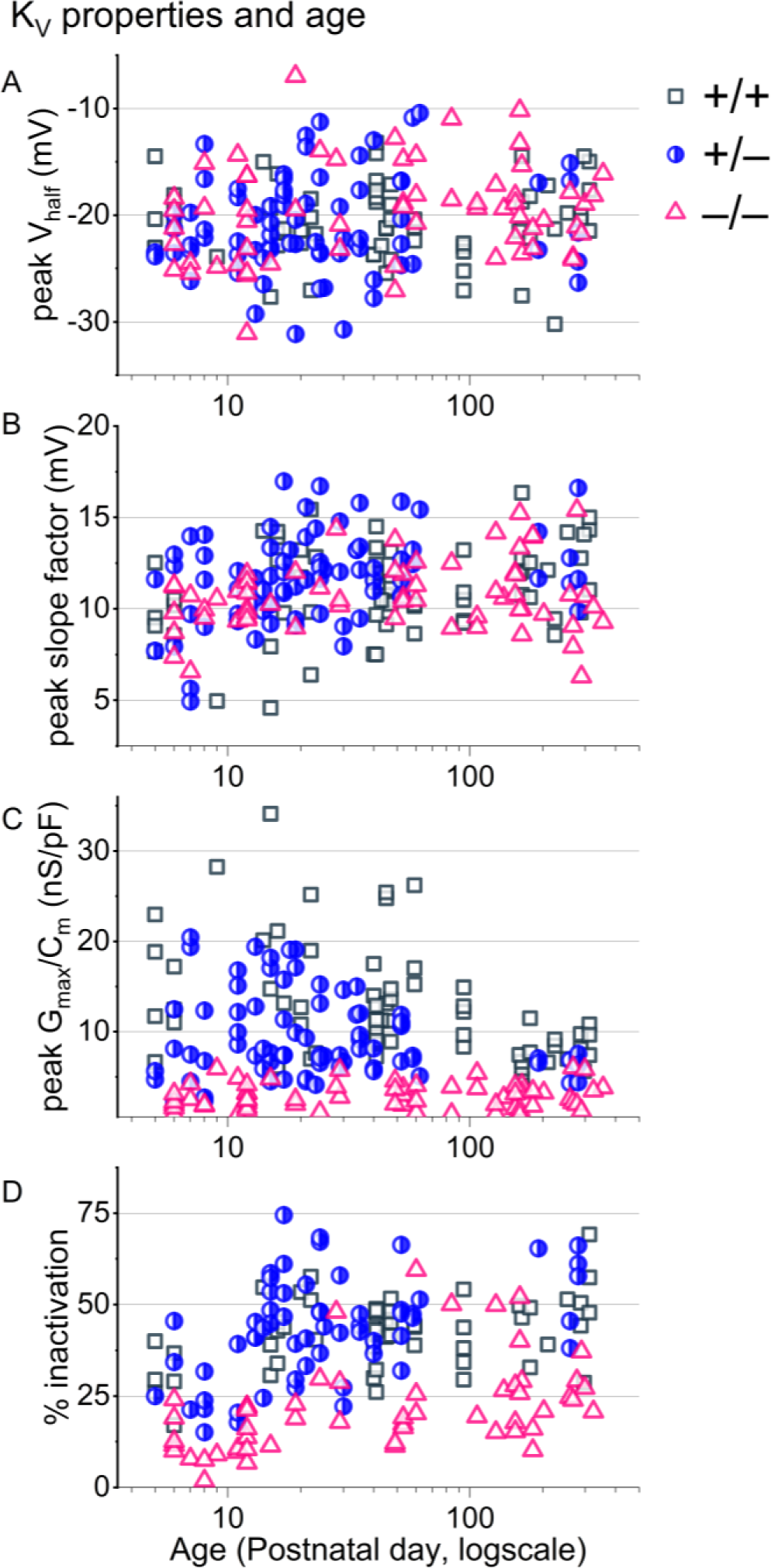
For type II HCs older than P12, K_V_ conductances were stable. (**A-C**) Parameters from Boltzmann fits of peak G-V relations and (**D**) % inactivation at +30 mV plotted against age from all zones. Overlaid curves are smoothing cubic β-splines. Note the seven extrastriolar K_V_1.8^−/−^ type II HCs with % inactivation >30%.

**Supplemental Figure 4.**
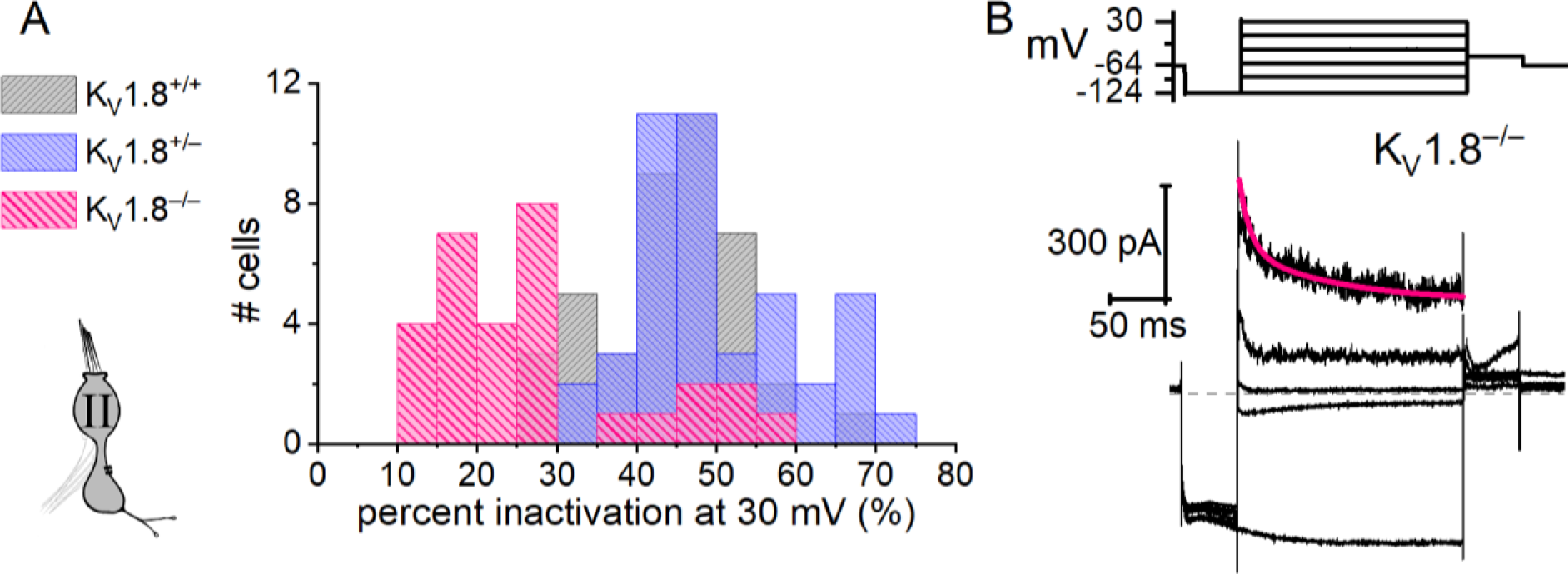
A minority of extrastriolar K_V_1.8^−/−^ type II HCs had a very small fast-inactivating outward rectifier current. (A) All extrastriolar K_V_1.8^+/+,+/−^ type II HCs inactivated by >30%. Most mature (>P12) extrastriolar K_V_1.8^−/−^ type II HCs inactivated by <30% but some inactivated by >30% (7/30, 23%) because they had fast inactivation (B). (B) Exemplar residual fast inactivation (τ_FastInact_ = 10 ms at +30 mV). For the 7 cells in this group, τ_FastInact_ = 30 ± 6 ms, amplitude of fast inactivation = 310 ± 70 pA; activation peak V_half_ = –15 ± 2 mV and slope factor = 12.4 ± 0.9 mV. These parameters are similar to g_A_ but for the much smaller conductance (one-way ANOVAs).

**Supplemental Figure 5.**
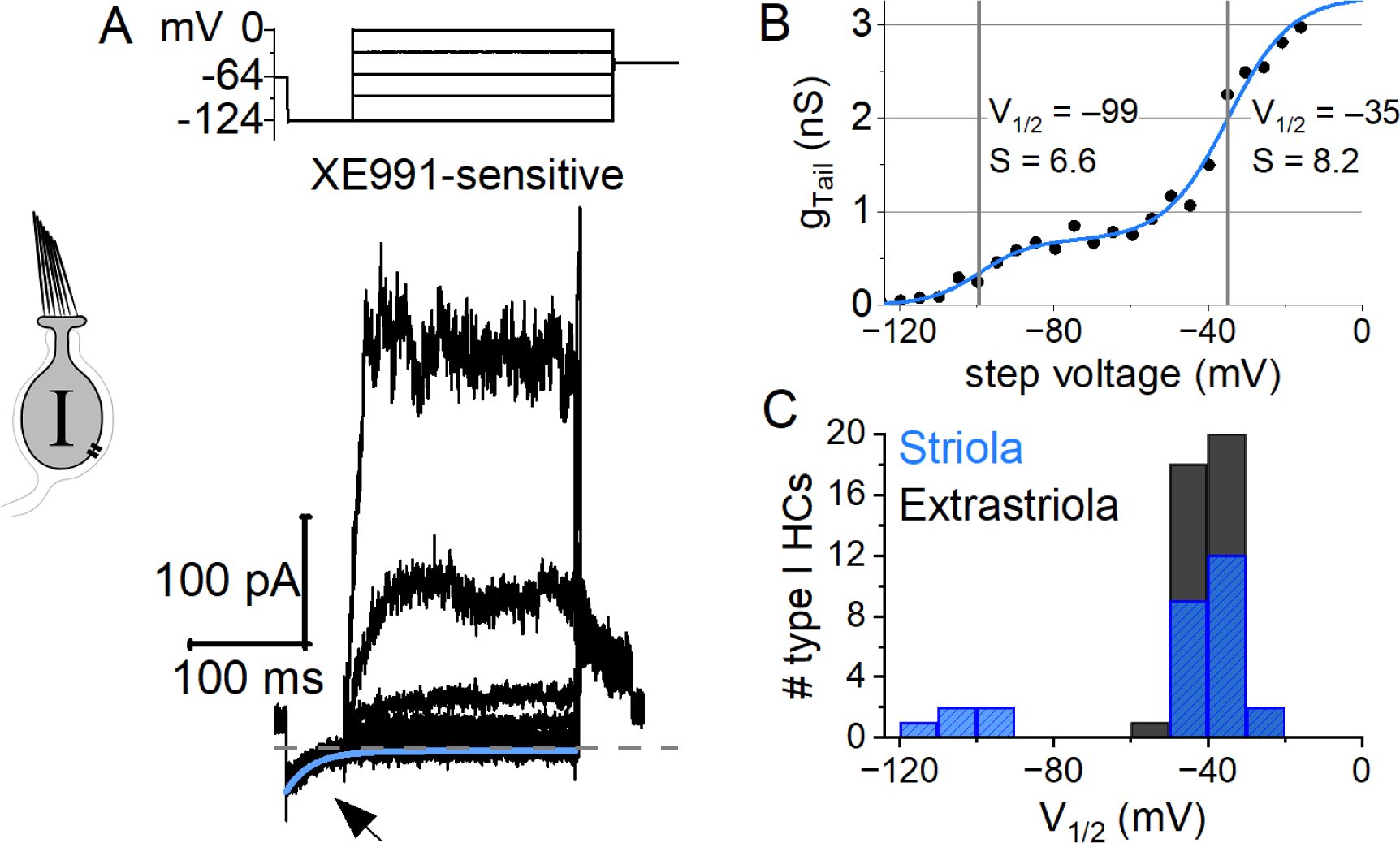
A minority of striolar K_V_1.8^−/−^ type I HCs had a small low-voltage-activated outward rectifier current in addition to a more positively activating outward rectifier. (**A**) Low-voltage-activated current from one cell was isolated by subtraction with 10 μM XE991 (P39), indicating that it was carried by K_V_7 channels. Deactivation of XE991-sensitive current evoked by step from –64 mV to –124 mV *(arrow*) was fit with exponential decay (τ = 21 ms). (**B**) XE991-sensitive tail G-V curve of the XE991-blocked conductance (A) was fit with a sum of two Boltzmann equations: G(V) = A_1_/(1 + exp((V_half,1_ – V)/S_1_)) + A_2_/(1 + exp((V_half,2_ – V)/S_2_)). (**C**) The low-voltage-activated V_half,1_ component was only seen in striolar K_V_1.8^−/−^ type I HCs, and even there in the minority: 5/23; 22%; P6-P370). It was always seen together with a more positively activating outward rectifier. Boltzmann parameters, including (B): A_1_/(A_1_+A_2_) = 0.15 ± 0.04, V_half,1_ = –106 ± 5 mV (n=5), S_1_ = 3.8 ± 0.8 mV, V_half,2_ = –41 ± 1 mV, S_2_ = 7 ± 1 mV. Ages: P11, 39, 202, 202, 202. No extrastriolar type I HCs (0/45; P6-277) had a double activation tail G-V curve.

**Supplemental Figure 6.**
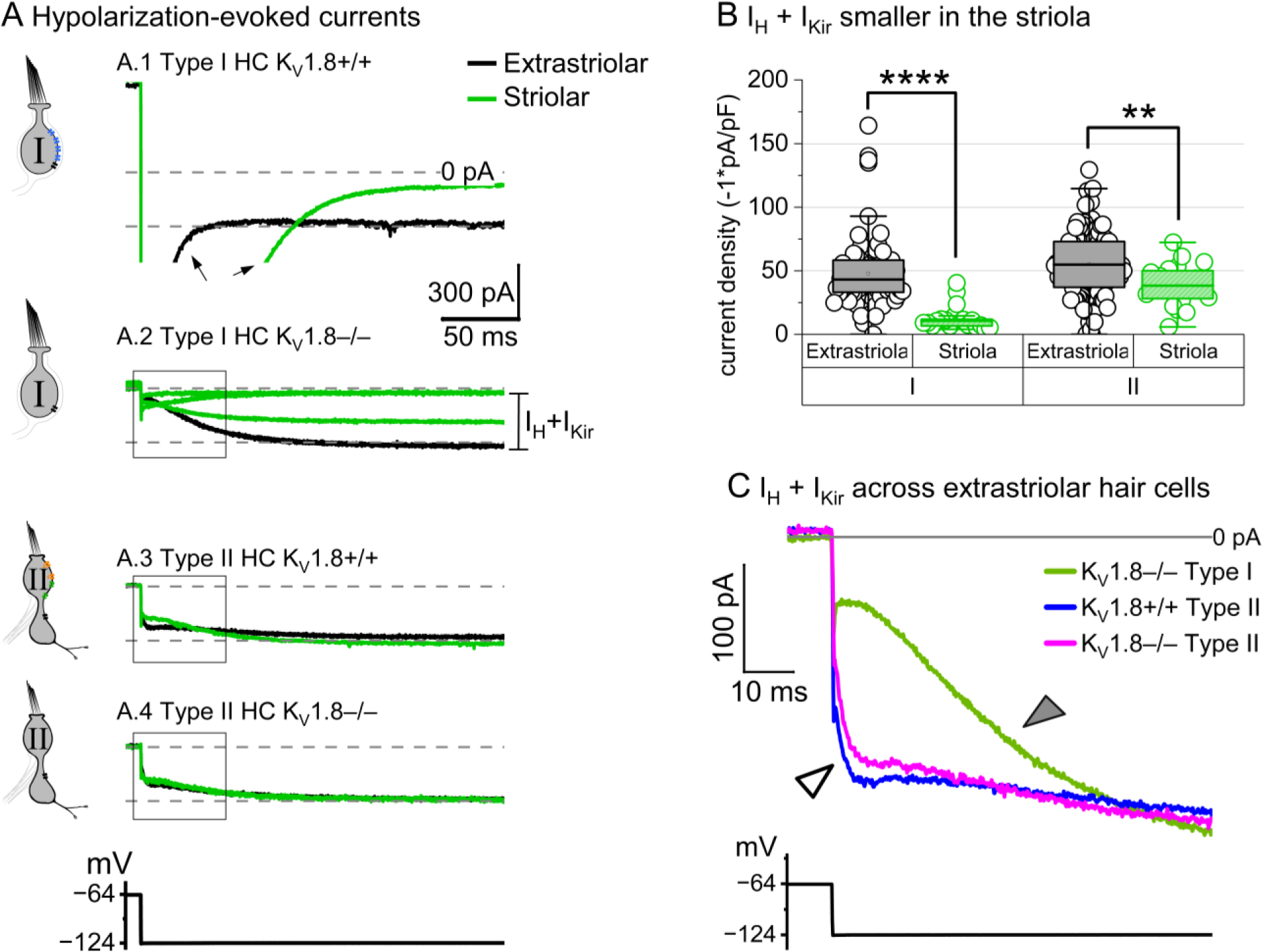
No difference was detected in H (HCN) and Kir (fast inward rectifier) currents between K_V_1.8^+/+^ and K_V_1.8^−/−^ hair cells, consistent with a specific involvement of K_V_1.8 in *Kcna10* expression. (A) Hyperpolarizing voltage steps evoked I_Kir_ and I_HCN_ in K_V_1.8^+/+,+/–,–/−^ type I and II HCs. **A.1** *Arrows*, deactivation of g_K,L_. **A.2-A.4** *Bracket,* the sum of I_H_ and I_Kir_ were measured as total inward current after 250 ms at –124 mV. (B) Summed I_KIR_ and I_H_ density was the same across genotypes but smaller in striola than extrastriola (see Supplemental Table 3 for statistics). (C) Inward currents seen at the onset of hyperpolarization, including I_Kir_, were larger in type II HCs than type I. Magnification of boxed in inset from the extrastriolar cells in A.2-A.4. *Open arrowhead*, activation of fast inward rectifier, I_Kir_; *filled arrowhead*, slower activation of I_HCN_.

**Supplemental Table 1.**
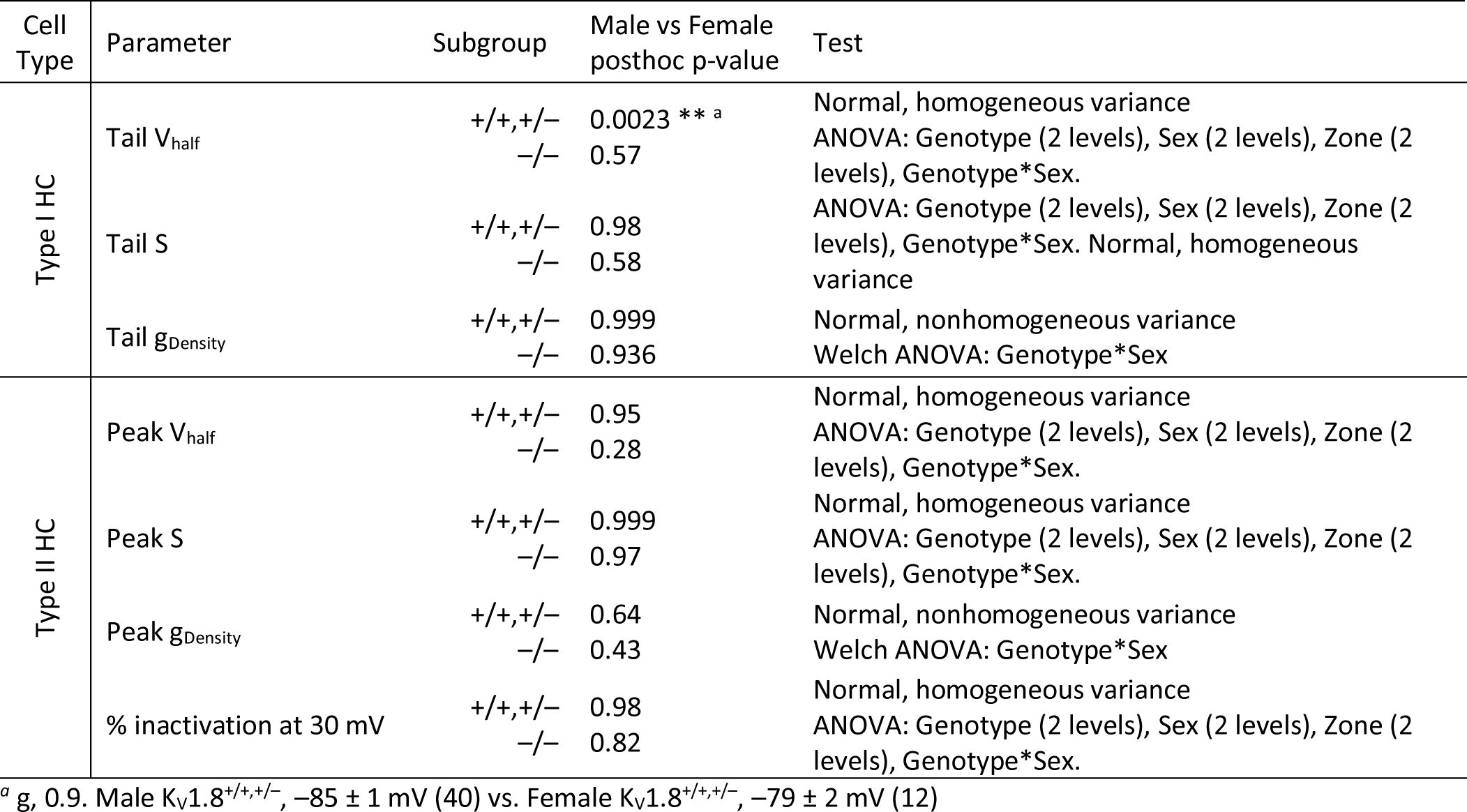
Test of sex differences in hair cell K_V_ channel data.

**Supplemental Table 2.** Soma size of type I HCs. The perimeter of cross-sections of basolateral cell bodies was manually measured from immunohistochemistry sections with sufficient autofluorescence in the 488 channel to distinguish cell morphology, and anti-calretinin to label type II hair cells and calyx-only afferents. Cells were measured from two female littermates.

| Cell Type | Genotype | Age | Weight (g) | Perimeter (μm) | Test |
| --- | --- | --- | --- | --- | --- |
| Type I HC | +/+ | P117 | 24.9 | 47 ± 1 (19) | ANOVA: p=0.1,<br>power = 0.07 |
| | -/- | P117 | 25.2 | 46 ± 1 (28) | |

**Supplemental Table 3.**
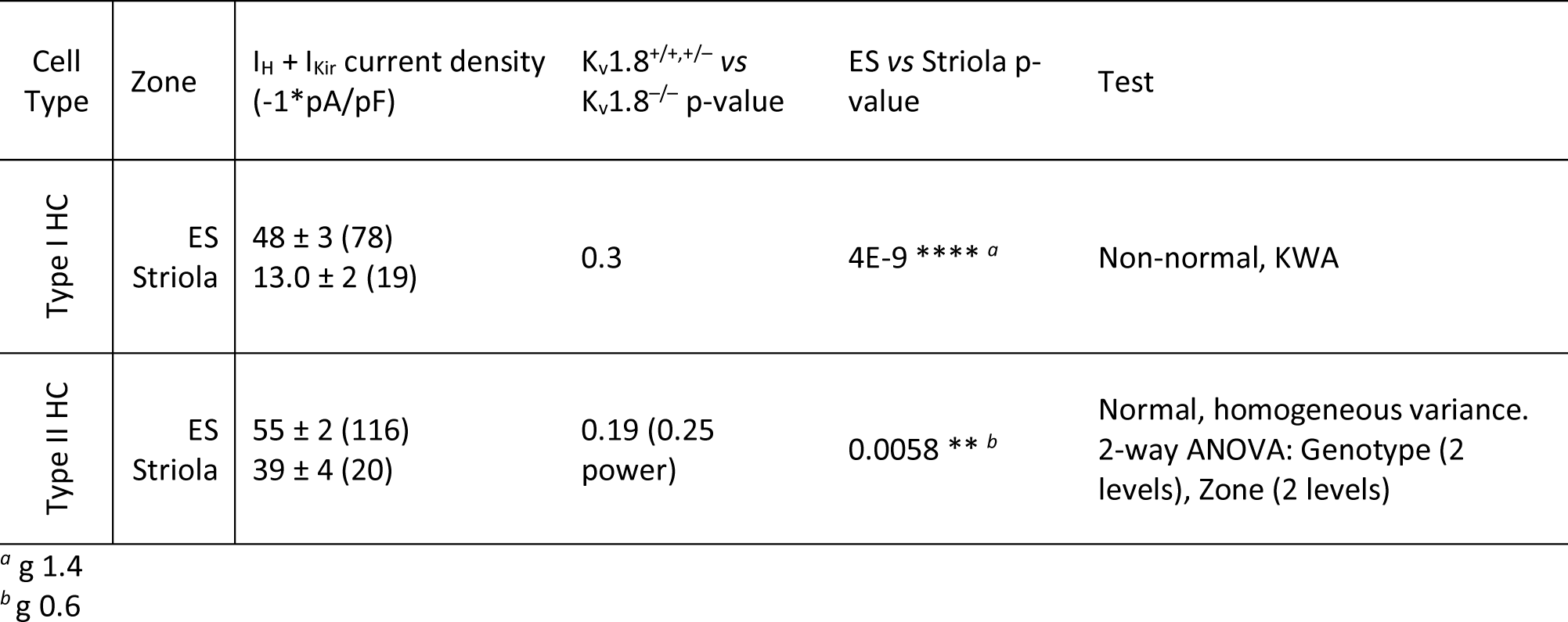
Detected zonal but not genotype differences in hair cell I_Kir_ and I_H_.

## References

Alexander SPH et al. (2019) The Concise Guide to Pharmacology 2019/20: Ion channels. Br J Pharmacol 176 Available at: https://onlinelibrary.wiley.com/doi/10.1111/bph.14749 [Accessed April 12, 2022].

Al-Sabi A, Kaza S, Le Berre M, O’Hara L, Bodeker M, Wang J, Dolly JO (2011) Position-dependent attenuation by Kv1.6 of N-type inactivation of Kv1.4-containing channels. Biochem J 438:389– 396.

Ashmore JF (1983) Frequency tuning in a frog vestibular organ. Nature 304:536–538.

Bao H, Wong WH, Goldberg JM, Eatock RA (2003) Voltage-Gated Calcium Channel Currents in Type I and Type II Hair Cells Isolated From the Rat Crista. J Neurophysiol 90:155–164.

Baukrowitz T, Yellen G (1995) Modulation of K+ current by frequency and external [K+]: A tale of two inactivation mechanisms. Neuron 15:951–960.

Behrend O, Schwark C, Kunihiro T, Strupp M (1997) Cyclic GMP inhibits and shifts the activation curve of the delayed rectifier (I(K1)) of type I mammalian vestibular hair cells. NeuroReport 8:2687–2690.

Brown DA, Selyanko AA, Hadley JK, Tatulian L (2002) Some Pharmacological Properties Of Neural KCNQ Channels. Neurophysiology 34:91–94.

Carlisle FA, Steel KP, Lewis MA (2012) Specific expression of Kcna10, Pxn and Odf2 in the organ of Corti. Gene Expression Patterns 12:172–179.

Chen JWY, Eatock RA (2000) Major Potassium Conductance in Type I Hair Cells from Rat Semicircular Canals: Characterization and Modulation by Nitric Oxide. J Neurophysiol 84:139–151.

Contini D, Holstein GR, Art JJ (2020) Synaptic cleft microenvironment influences potassium permeation and synaptic transmission in hair cells surrounded by calyx afferents in the turtle. J Physiol 598:853–889.

Contini D, Price SD, Art JJ (2017) Accumulation of K^+^ in the synaptic cleft modulates activity by influencing both vestibular hair cell and calyx afferent in the turtle: K^+^ modulation of synaptic transmission between hair cell and afferent. J Physiol 595:777–803.

Contini D, Zampini V, Tavazzani E, Magistretti J, Russo G, Prigioni I, Masetto S (2012) Intercellular K^+^ accumulation depolarizes Type I vestibular hair cells and their associated afferent nerve calyx. Neuroscience 227:232–246.

Correia MJ, Lang DG (1990) An electrophysiological comparison of solitary type I and type II vestibular hair cells. Neuroscience Letters 116:106–111.

Correia MJ, Ricci AJ, Rennie KJ (1996) Filtering Properties of Vestibular Hair Cells: An Update. Ann NY Acad Sci 781:138–149.

Correia MJ, Weng T, Prusak D, Wood TG (2008) Kvβ1.1 Associates with Kv1.4 in Chinese Hamster Ovary Cells and Pigeon Type II Vestibular Hair Cells and Enhances the Amplitude, Inactivation and Negatively Shifts the Steady-State Inactivation Range. Neuroscience 152:809–820.

Crawford AC, Fettiplace R (1981) An electrical tuning mechanism in turtle cochlear hair cells. J Physiol 312:377–412.

Desai SS, Zeh C, Lysakowski A (2005) Comparative Morphology of Rodent Vestibular Periphery. I. Saccular and Utricular Maculae. J Neurophysiol 93:251–266.

Dierich M, Altoè A, Koppelmann J, Evers S, Renigunta V, Schäfer MK, Naumann R, Verhulst S, Oliver D, Leitner MG (2020) Optimized Tuning of Auditory Inner Hair Cells to Encode Complex Sound through Synergistic Activity of Six Independent K+ Current Entities. Cell Reports 32:107869.

Dwenger MM, Raph SM, Baba SP, Moore JB, Nystoriak MA (2022) Diversification of Potassium Currents in Excitable Cells via Kvβ Proteins. Cells 11:2230.

Eatock RA, Songer JE (2011) Vestibular Hair Cells and Afferents: Two Channels for Head Motion Signals. Annu Rev Neurosci 34:501–534.

Erickson T, Nicolson T (2015) Identification of sensory hair-cell transcripts by thiouracil-tagging in zebrafish. BMC Genomics 16:842.

Fettiplace R (1987) Electrical tuning of hair cells in the inner ear. TINS 10:5.

Géléoc GS, Risner J, Holt JR (2004) Developmental Acquisition of Voltage-Dependent Conductances and Sensory Signaling in Hair Cells of the Embryonic Mouse Inner Ear. J Neurosci 24:11148–11159.

Goldberg JM (2000) Afferent diversity and the organization of central vestibular pathways. Exp Brain Res 130:277–297.

González-Garrido A, Pujol R, López-Ramírez O, Finkbeiner C, Eatock RA, Stone JS (2021) The Differentiation Status of Hair Cells That Regenerate Naturally in the Vestibular Inner Ear of the Adult Mouse. J Neurosci 41:7779–7796.

Govindaraju AC, Quraishi IH, Lysakowski A, Eatock RA, Raphael RM (2023) Nonquantal transmission at the vestibular hair cell–calyx synapse: K_LV_ currents modulate fast electrical and slow K ^+^ potentials. Proc Natl Acad Sci USA 120:e2207466120.

Gulley A, Bagger-Sjöbäck D (1979) Freeze-fracture studies on the synapse between the type I hair cells and the calyceal terminal in the guinea-pig vestibular system. J Neurocytol 8:591–603.

Holt JR, Stauffer EA, Abraham D, Geleoc GSG (2007) Dominant-Negative Inhibition of M-Like Potassium Conductances in Hair Cells of the Mouse Inner Ear. J Neurosci 27:8940–8951.

Horwitz GC, Risner-Janiczek JR, Jones SM, Holt JR (2011) HCN Channels Expressed in the Inner Ear Are Necessary for Normal Balance Function. J Neurosci 31:16814–16825.

Hudspeth AJ, Lewis RS (1988) A model for electrical resonance and frequency tuning in saccular hair cells of the bull-frog, Rana catesbeiana. J Physiol 400:275–297.

Hurley KM, Gaboyard S, Zhong M, Price SD, Wooltorton JRA, Lysakowski A, Eatock RA (2006) M-Like K^+^ Currents in Type I Hair Cells and Calyx Afferent Endings of the Developing Rat Utricle. Journal of Neuroscience 26:10253–10269.

Imbrici P, D’Adamo MC, Kullmann DM, Pessia M (2006) Episodic ataxia type 1 mutations in the *KCNA1* gene impair the fast inactivation properties of the human potassium channels Kv1.4-1.1/Kvβ1.1 and Kv1.4-1.1/Kvβ1.2. Eur J Neurosci 24:3073–3083.

Jan TA, Eltawil Y, Ling AH, Chen L, Ellwanger DC, Heller S, Cheng AG (2021) Spatiotemporal dynamics of inner ear sensory and non-sensory cells revealed by single-cell transcriptomics. Cell Reports 36:109358.

Jensen HS, Grunnet M, Olesen S-P (2007) Inactivation as a New Regulatory Mechanism for Neuronal Kv7 Channels. Biophys J 92:2747–2756.

Kharkovets T, Hardelin J-P, Safieddine S, Schweizer M, El-Amraoui A, Petit C, Jentsch TJ (2000) KCNQ4, a K^+^ channel mutated in a form of dominant deafness, is expressed in the inner ear and the central auditory pathway. Proc Natl Acad Sci USA 97:4333–4338.

Kole MJ, Qian J, Waase MP, Klassen TL, Chen TT, Augustine GJ, Noebels JL (2015) Selective Loss of Presynaptic Potassium Channel Clusters at the Cerebellar Basket Cell Terminal Pinceau in Adam11 Mutants Reveals Their Role in Ephaptic Control of Purkinje Cell Firing. J Neurosci 35:11433–11444.

Kubisch C, Schroeder BC, Friedrich T, Lütjohann B, El-Amraoui A, Marlin S, Petit C, Jentsch TJ (1999) KCNQ4, a Novel Potassium Channel Expressed in Sensory Outer Hair Cells, Is Mutated in Dominant Deafness. Cell 96:437–446.

Lang R, Lee G, Liu W, Tian S, Rafi H, Orias M, Segal AS, Desir GV (2000) KCNA10: A novel ion channel functionally related to both voltage-gated potassium and CNG cation channels. American Journal of Physiology - Renal Physiology 278.

Lee SI, Conrad T, Jones SM, Lagziel A, Starost MF, Belyantseva IA, Friedman TB, Morell RJ (2013) A null mutation of mouse Kcna10 causes significant vestibular and mild hearing dysfunction. Hear Res 300:1–9.

Levin ME, Holt JR (2012) The function and molecular identity of inward rectifier channels in vestibular hair cells of the mouse inner ear. J Neurophysiol 108:175–186.

Lewis ER (1988) Tuning in the bullfrog ear. Biophysical Journal 53:441–447.

Lim R, Kindig AE, Donne SW, Callister RJ, Brichta AM (2011) Potassium accumulation between type I hair cells and calyx terminals in mouse crista. Exp Brain Res 210:607–621.

Liu H, Chen L, Giffen KP, Stringham ST, Li Y, Judge PD, Beisel KW, He DZZ (2018) Cell-Specific Transcriptome Analysis Shows That Adult Pillar and Deiters’ Cells Express Genes Encoding Machinery for Specializations of Cochlear Hair Cells. Front Mol Neurosci 11:356.

López-Barneo J, Hoshi T, Heinemann SH, Aldrich RW (1993) Effects of external cations and mutations in the pore region on C-type inactivation of Shaker potassium channels. Recept Channels 1:61–71.

Lysakowski A, Gaboyard-Niay S, Calin-Jageman I, Chatlani S, Price SD, Eatock RA (2011) Molecular Microdomains in a Sensory Terminal, the Vestibular Calyx Ending. J Neurosci 31:10101–10114.

Lysakowski A, Goldberg JM (2004) Morphophysiology of the vestibular periphery. In: The Vestibular System, pp 57–152 Springer Hearing and Auditory Research. Springer New York. Available at: 10.1007/0-387-21567-0_3.

Marino M, Misuri L, Brogioli D (2014) A new open source software for the calculation of the liquid junction potential between two solutions according to the stationary Nernst-Planck equation. Available at: http://arxiv.org/abs/1403.3640 [Accessed October 23, 2023].

Masetto S, Correia MJ (1997) Electrophysiological Properties of Vestibular Sensory and Supporting Cells in the Labyrinth Slice Before and During Regeneration. J Neurophysiol 78:1913–1927.

McInturff S, Burns JC, Kelley MW (2018) Characterization of spatial and temporal development of Type I and Type II hair cells in the mouse utricle using new cell-type-specific markers. Biol Open 7:bio038083.

Meredith FL, Rennie KJ (2016) Channeling your inner ear potassium: K+ channels in vestibular hair cells. Hear Res 338:40–51.

Mitra K, Ubarretxena-Belandia I, Taguchi T, Warren G, Engelman DM (2004) Modulation of the bilayer thickness of exocytic pathway membranes by membrane proteins rather than cholesterol. Proc Natl Acad Sci USA 101:4083–4088.

Orvis J et al. (2021) gEAR: Gene Expression Analysis Resource portal for community-driven, multi-omic data exploration. Nat Methods 18:843–844.

Pastras CJ, Curthoys IS, Asadnia M, McAlpine D, Rabbitt RD, Brown DJ (2023) Evidence that ultrafast non-quantal transmission underlies synchronized vestibular action potential generation. J Neurosci:JN-RM-1417–23.

Perez-Flores MC, Lee JH, Park S, Zhang X-D, Sihn C-R, Ledford HA, Wang W, Kim HJ, Timofeyev V, Yarov-Yarovoy V, Chiamvimonvat N, Rabbitt RD, Yamoah EN (2020) Cooperativity of K _v_ 7.4 channels confers ultrafast electromechanical sensitivity and emergent properties in cochlear outer hair cells. Sci Adv 6:eaba1104.

Pujol R, Pickett SB, Nguyen TB, Stone JS (2014) Large basolateral processes on type II hair cells are novel processing units in mammalian vestibular organs: Basolateral Processes of Type II Hair Cells. J Comp Neurol 522:3141–3159.

Ramanathan K, Fuchs PA (2002) Modeling Hair Cell Tuning by Expression Gradients of Potassium Channel. Biophys J 82:64–72.

Ranjan R, Logette E, Marani M, Herzog M, Tache V, Scantamburlo E, Buchilier V, Markram H (2019) A Kinetic Map of the Homomeric Voltage-Gated Potassium Channel (Kv) Family. Front Cell Neurosci 13:358.

Rennie KJ, Correia MJ (1994) Potassium currents in mammalian and avian isolated type I semicircular canal hair cells. J Neurophysiol 71:317–329.

Rennie KJ, Weng T, Correia MJ (2001) Effects of KCNQ channel blockers on K^+^ currents in vestibular hair cells. Am J Physiol Cell Physiol 280:C473–C480.

Ricci AJ, Rennie KJ, Correia MJ (1996) The delayed rectifier, IKI, is the major conductance in type I vestibular hair cells across vestibular end organs. Pflüg Arch Eur J Physiol 432:34–42.

Rothman JS, Manis PB (2003) Kinetic Analyses of Three Distinct Potassium Conductances in Ventral Cochlear Nucleus Neurons. J Neurophysiol 89:3083–3096.

Rüsch A, Eatock RA (1996a) A delayed rectifier conductance in type I hair cells of the mouse utricle. J Neurophysiol 76:995–1004.

Rüsch A, Eatock RA (1996b) Voltage Responses of Mouse Utricular Hair Cells to Injected Currents. Ann NY Acad Sci 781:71–84.

Rüsch A, Lysakowski A, Eatock RA (1998) Postnatal development of type I and type II hair cells in the mouse utricle: Acquisition of voltage-gated conductances and differentiated morphology. Journal of Neuroscience 18:7487–7501.

Scheffer DI, Shen J, Corey DP, Chen Z-Y (2015) Gene Expression by Mouse Inner Ear Hair Cells during Development. J Neurosci 35:6366–6380.

Scheibinger M, Janesick A, Benkafadar N, Ellwanger DC, Jan TA, Heller S (2022) Cell-type identity of the avian utricle. Cell Reports 40:111432.

Schroeder BC, Hechenberger M, Weinreich F, Kubisch C, Jentsch TJ (2000) KCNQ5, a Novel Potassium Channel Broadly Expressed in Brain, Mediates M-type Currents. Journal of Biological Chemistry 275:24089–24095.

Schweizer FE, Savin D, Luu C, Sultemeier DR, Hoffman LF (2009) Distribution of high-conductance calcium-activated potassium channels in rat vestibular epithelia. J Comp Neurol 517:134–145.

Smith GT, Proffitt MR, Smith AR, Rusch DB (2018) Genes linked to species diversity in a sexually dimorphic communication signal in electric fish. J Comp Physiol A 204:93–112.

Songer JE, Eatock RA (2013) Tuning and Timing in Mammalian Type I Hair Cells and Calyceal Synapses. J Neurosci 33:3706–3724.

Sousa AD, Andrade LR, Salles FT, Pillai AM, Buttermore ED, Bhat MA, Kachar B (2009) The Septate Junction Protein Caspr Is Required for Structural Support and Retention of KCNQ4 at Calyceal Synapses of Vestibular Hair Cells. J Neurosci 29:3103–3108.

Spaiardi P, Tavazzani E, Manca M, Milesi V, Russo G, Prigioni I, Marcotti W, Magistretti J, Masetto S (2017) An allosteric gating model recapitulates the biophysical properties of *I* _K,L_ expressed in mouse vestibular type I hair cells: Allosteric gating of *I* _K,L_. J Physiol 595:6735–6750.

Spaiardi P, Tavazzani E, Manca M, Russo G, Prigioni I, Biella G, Giunta R, Johnson SL, Marcotti W, Masetto S (2020) K+ Accumulation and Clearance in the Calyx Synaptic Cleft of Type I Mouse Vestibular Hair Cells. Neuroscience 426:69–86.

Spitzmaul G, Tolosa L, Winkelman BHJ, Heidenreich M, Frens MA, Chabbert C, de Zeeuw CI, Jentsch TJ (2013) Vestibular Role of KCNQ4 and KCNQ5 K+ Channels Revealed by Mouse Models*. Journal of Biological Chemistry 288:9334–9344.

Stone JS, Pujol R, Nguyen TB, Cox BC (2021) The Transcription Factor Sox2 Is Required to Maintain the Cell Type-Specific Properties and Innervation of Type II Vestibular Hair Cells in Adult Mice. J Neurosci 41:6217–6233.

Stühmer W, Ruppersberg JP, Schröter KH, Sakmann B, Stocker M, Giese KP, Perschke A, Baumann A, Pongs O (1989) Molecular basis of functional diversity of voltage-gated potassium channels in mammalian brain. EMBO J 8:3235–3244.

Vollrath MA, Eatock RA (2003) Time course and extent of mechanotransducer adaptation in mouse utricular hair cells: Comparison with frog saccular hair cells. Journal of Neurophysiology 90:2676–2689.

Wang H (1998) KCNQ2 and KCNQ3 Potassium Channel Subunits: Molecular Correlates of the M-Channel. Science 282:1890–1893.

Wersäll J (1956) Studies on the structure and innervation of the sensory epithelium of the cristae ampulares in the guinea pig; a light and electron microscopic investigation. Acta Otolaryngol Suppl 126:1–85.

Wong WH, Hurley KM, Eatock RA (2004) Differences Between the Negatively Activating Potassium Conductances of Mammalian Cochlear and Vestibular Hair Cells. J Assoc Res Otolaryngol 5:270– 284.

Xu T, Nie L, Zhang Y, Mo J, Feng W, Wei D, Petrov E, Calisto LE, Kachar B, Beisel KW, Vazquez AE, Yamoah EN (2007) Roles of Alternative Splicing in the Functional Properties of Inner Ear-specific KCNQ4 Channels. J Biol Chem 282:23899–23909.

Yao X (2002) Expression of KCNA10, a Voltage-Gated K Channel, in Glomerular Endothelium and at the Apical Membrane of the Renal Proximal Tubule. Journal of the American Society of Nephrology 13:2831–2839.

